# Cas12a/3 crRNAs RNP transformation enables transgene-free multiplex genome editing, long deletions, and inversions in citrus chromosome in the T0 generation

**DOI:** 10.1101/2024.06.13.598908

**Authors:** Hang Su, Yuanchun Wang, Jin Xu, Ahmad A. Omar, Jude W. Grosser, Nian Wang

**Author notes:** These authors contributed equally: Hang Su, Yuanchun Wang. Correspondence to Nian Wang.

## Abstract

Citrus canker, caused by *Xanthomonas citri* subsp. citri (*Xcc*), is a devastating disease worldwide. Previously, we successfully generated canker-resistant *Citrus sinensis* cv. Hamlin lines in the T0 generation, achieving a mutation efficiency of 97.4%. This was achieved through the transformation of embryogenic protoplasts using the Cas12a/1 crRNA ribonucleoprotein (RNP) system to edit the canker susceptibility gene, *CsLOB1*, which led to small indels. Here, we transformed embryogenic protoplasts of Hamlin with Cas12a/3 crRNAs RNP, resulting in 100% efficiency in editing the *CsLOB1* gene in the T0 generation. Among the 10 transgene-free genome-edited lines, long deletions were obtained in five lines. Additionally, inversions were observed in three of the five edited lines with long deletions, but not in any edited lines with short indel mutations, suggesting long deletions are required for inversions. Biallelic mutations were observed for each of the three target sites in 4 of the 10 edited lines when 3 crRNAs were used, demonstrating that transformation of embryogenic citrus protoplasts with Cas12a/3 crRNAs RNP can be very efficient for multiplex editing. Our analysis revealed the absence of off-target mutations in the edited lines. These *cslob1* mutant lines were canker-resistant and no canker symptoms were observed after inoculation with *Xcc* and *Xcc* growth was significantly reduced in the *cslob1* mutant lines compared to the wild type plants. Taken together, Cas12a/3 crRNAs RNP transformation of embryogenic protoplasts of citrus provides a promising solution for transgene-free multiplex genome editing with high efficiency and for deletion of long fragments.

## Introduction

CRISPR/Cas mediated genome editing has shown enormous potential in genetic improvements of crops. It has been used to improve disease resistance, yield, quality, and tolerance against abiotic stress among others (Zhu et al., 2020). For the adoption of the genome-edited crops, the plants are usually required to be transgene-free to address the public concerns and regulatory issues (Turnbull et al., 2021). In addition, other putative impacts have also been reported to result from transgenic expression of CRISPR/Cas including unintended genome editing owing to the consistent expression of CRISPR/Cas and disruption of gene functions at the insertion site (O’Malley and Ecker, 2010; He et al., 2018; Sturme et al., 2022). Most genome-edited plants were generated via *Agrobacterium*-mediated transformation resulting in transgenic plants. For the transgenic genome-edited plants, foreign genes can be removed relatively easily for annual crops by segregation through selfing or backcrossing T0 plants to wild-type plants (Bhattacharjee et al., 2023) and using a transgene-killer strategy (He et al., 2018). In addition, multiple technologies have been reported to generate transgene-free genome-edited plants in the T0 generation. For instance, transient expression of ribonucleoproteins of Cas9/gRNA or Cas12a/crRNA, as well as CRISPR/Cas DNA and RNA have been successfully used to generate transgene-free genome edited *Arabidopsis thaliana*, tobacco, lettuce, rice, wheat grape, apple, soybean, and cabbage (Woo et al., 2015; Malnoy et al., 2016; Svitashev et al., 2016; Zhang et al., 2016; Kim et al., 2017; Liang et al., 2017; Murovec et al., 2018). Viral vectors have also been used to generate transgene-free plants (Ma et al., 2020).

Citrus is one of the top three tree crops worldwide. Traditional citrus breeding usually takes 20-30 years owing to the long juvenility (Gmitter et al., 2012). Cas9/sgRNA was firstly used for citrus genome editing in 2013 (Jia and Wang, 2014a). Significant progress in citrus genome editing has since been made (Jia and Wang, 2014b; Jia et al., 2016; Jia et al., 2017; LeBlanc et al., 2017; Peng et al., 2017; Zhang et al., 2017b; Jia et al., 2019; Zhu et al., 2019; Zou et al., 2019; Dutt et al., 2020; Huang et al., 2020a; Huang et al., 2021; Jia et al., 2022; Mahmoud et al., 2022). It was reported that the biallelic/homozygous mutation rates was 89% for Carrizo citrange and 79% for Hamlin sweet orange via *Agrobacterium*-mediated transformation of epicotyls (Huang et al., 2021). Interestingly, genome editing via hairy root genetic transformation using *Agrobacterium rhizogenes* has also shown promises (Irigoyen et al., 2020; Ma et al., 2022; Wang et al., 2023).

CRISPR/Cas genome editing has successfully improved disease resistance against citrus canker by mutating *LOB1*, the canker susceptibility gene (Hu et al., 2014; Jia et al., 2016; Jia et al., 2017; Peng et al., 2017; Huang et al., 2021). Citrus canker caused by *Xanthomonas citri* pv. citri (*Xcc*) is arguably one of the top two citrus diseases in the world (Gottwald et al., 2002; Ference et al., 2018). The *LOB1*-edited pummelo, grapefruit, and sweet orange all showed no canker symptoms when inoculated with *Xcc* (Jia et al., 2016; Jia et al., 2017; Peng et al., 2017; Huang et al., 2021;2022; Jia et al., 2022). Importantly, we have generated transgene-free canker resistant *C. sinensis* cv. Hamlin through transformation of embryogenic protoplasts with Cas12a/crRNA RNP (Su et al., 2023). In addition, a co-editing strategy based on *Agrobacterium* transformation has also been successfully used to generate transgene-free genome edited pummelo (Huang et al., 2023) and sweet orange (Jia et al. 2024).

In a previous study, one crRNA was used to guide the genome editing of the canker susceptibility gene *LOB1* to generate transgene-free canker-resistant *C. sinensis* cv. Hamlin (Su et al., 2023). The mutations were mostly short deletions. Multiple gRNAs have been used to improve genome editing efficiency and generate long deletions (Xie et al., 2015). Here, we have conducted genome editing of the *LOB1* gene in *C. sinensis* cv. Hamlin using 3 crRNAs through transformation of embryogenic protoplasts with Cas12a/3 crRNAs RNP. Three crRNAs indeed achieved high genome editing efficiency, led to multiplex genome editing, long deletions as well as inversions.

## Materials and methods

### Citrus plants and cell culture conditions

*C. sinensis* seedlings were grown in a greenhouse at the Citrus Research and Education Center in Lake Alfred, Florida. Embryogenic callus lines originating from immature ovules of *C. sinensis* cv. Hamlin were established and maintained using the Murashige and Tucker (1969, MT) medium (PhytoTech Labs, Lenexa, KS, USA). This medium was supplemented with 5.0 mg/L of kinetin (KIN) and 500 mg/L of malt extract. Meanwhile, the suspension cell culture of *C. sinensis* cv. Hamlin was maintained in darkness at 22°C and sub-cultured biweekly. The cultivation medium was the H+H medium (Omar et al., 2016a). After 7-10 days following subculturing, the suspension cells were used for protoplast isolation.

### Protoplast isolation

Protoplast isolation was conducted as described previously (Su et al., 2023). Briefly, embryogenic *C. sinensis* cv. Hamlin protoplasts were isolated from the suspension cells with digestion solution (2.5 × volume BH3 (Omar et al., 2016a) and 1.5× volume enzyme solution (0.7 M mannitol, 24 mM CaCl_2_, 6.15 mM MES buffer, 2.4 % (w/v) Cellulase Onozuka RS (MX7353, Yakult Honsha, Minato-ku, Tokyo, Japan), 2.4 % (w/v) Macerozyme R-10 (MX7351, Yakult Honsha), pH 5.6) for 16-20 hours at 28 °C. The digested protoplast mixture was then filtered with a 40 μm cell strainer (431750, Corning, Durham, NC, USA) into a 50 mL Falcon tube, which were centrifuged at 60 g for 7 min. The pellets were resuspended with BH3 medium to rinse the protoplast. After repeating the washing step, the protoplasts were resuspended in 2 mL BH3 medium and diluted to 1 ×10^6^ cell/mL and kept in dark at room temperature for 1 hour.

### Transformation of embryogenic citrus protoplasts and plant regeneration

Protoplast transformation with RNP and plant regeneration were conducted as previously described (Omar et al., 2016a; Liu et al., 2023). Briefly, for RNP assembly, 0.81 nmol LbCas12U protein and 0.45 nmol of each of three crRNAs were assembled in 1 x Nuclease Reaction Buffer (NEB). Protein and crRNAs were mixed and incubated for 10 min at room temperature. For transfection, 1 mL protoplast cells, preassembled RNP, and 1 mL PEG-CaCl_2_ (0.4 M mannitol, 100 mM CaCl_2_, and 40% PEG-4000) were mixed and kept at room temperature for 15 min in dark followed by washing with BH3 medium twice. Edited citrus protoplast cells were kept in liquid medium (1:1:1 (v:v:v) mixture of BH3 and EME sucrose 0.6 M and EME sucrose 0.15 M) for 3 – 4 weeks at 28 °C in dark without shaking. Then citrus cells were transferred to EME sucrose medium supplemented with 1:2 mixture of BH3 and EME maltose 0.15 M and kept at 28 °C for 3–4 weeks in dark. Calli were regenerated from protoplasts and transferred to EME maltose solid medium supplemented with 1:2 mixture of BH3 and EME maltose 0.15 M and kept at 28 °C in dark for 3 – 4 weeks to generate embryos. Embryos were transferred to EME maltose solid medium and kept at room temperature under light for 3 – 4 weeks. Embryos were transferred to solid EME1500 medium and kept at room temperature under light for 3 – 4 months to generate shoots. Small plantlets were transferred to MS medium and kept at room temperature for 3 – 4 weeks. The regenerated shoots were micro-grafted onto Carrizo citrange rootstock in liquid rooting media and kept in tissue culture room at 25 °C under light for 3 – 4 weeks, grown in stonewool cubes in a growth chamber at 25 °C under light for 1 month, then planted in soil.

### Mutation identification and off-target test

Genomic DNA was isolated from leaves of both wild-type and *cslob1-*edited lines of *C. sinensis* cv. Hamlin. The Table S1 lists the primers employed in the PCR. For PCR amplification, the CloneAmp HiFi PCR Premix (639298, TakaraBio USA, San Jose, CA, USA) was utilized, in accordance with the manufacturer’s guidelines. The amplification procedure consisted of an initial step at 98 °C for 30 seconds, followed by 40 cycles at 98 °C for 10 seconds, 54 °C for 10 seconds, and 72 °C for 45 seconds. A final extension was performed at 72 °C for 5 minutes. The resulting PCR amplicons were cloned and subjected to sequencing employing the amplifying primers. The cloning was conducted using the Zero Blunt TOPO PCR Cloning Kit (450245, Thermo Fisher, San Jose, CA, USA), followed by transformation into Stellar Competent Cells from Takara. For the amplification and Sanger sequencing of single colonies, M13-F (GTAAAACGACGGCCAGTG) and M13-R (CAGGAAACAGCTATGACC) primers were used.

The off-target candidate sites sequences were listed in Table S1. The off-target sites were predicted using CRISPR-P 2.0 program(Liu et al., 2017). Then the predicted off-target sites with TTTV PAM site were further confirmed as off-target candidate sites and aligning target sequence with whole genome using BLAST program. Based on the whole genome sequencing mapping results, mutations of off-target sites were detected using the SAMtools package version 1.2 and deepvariant program version 1.4.0. There were no any off-target mutations detected in mutated plants.

### DNA Library construction, sequencing, and data analysis

Following the manufacturer’s protocol of short read DNA sequencing from Illumina, the library was prepared. 150-bp paired-end reads were generated using the Illumina NovaSeq 6000 platform according to the manufacturer’s instructions at Novogene, China. The raw paired-end reads were filtered to remove low-quality reads using fastp program version 0.22.0(Chen et al., 2018b). On average, more than 17.21 Gb of high-quality data was generated for each edited sweet orange lines (Table S2). To identify the mutations (single nucleotide polymorphisms, deletions and insertions) in the edited plant genomes, high quality paired-end short genomic reads were mapped to sweet orange(Wang et al., 2021) reference genome using Bowtie2 software version 2.2.6(Langmead and Salzberg, 2012). Based on the mapping results, mutations were identified using the SAMtools package version 1.2(Li et al., 2009) and deepvariant program version 1.4.0(Poplin et al., 2018) The identified mutations were filtered by quality and sequence depth (mapping quality > 10 and mapping depth >10). The mutations of target sites were visualized using IGV software version 2.15.4(Robinson et al., 2011).

### Quantitative reverse-transcription PCR (qRT-PCR)

*Xcc* strain 306 was infiltrated into wild type *C. sinensis* cv. Hamlin and transgene-free *cslob1* mutants at the concentration of 1 × 10^7^ cfu/mL. The infiltration areas of the leaf samples were collected at 9 days post-inoculation (dpi) for RNA isolation. Four biological replicates were used with one leaf as one biological replicate. Total RNA was extracted by TRIzol Reagent (15596026, Thermo-Fisher) following the manufacturer’s instructions. cDNA was synthesized by qScript cDNA SuperMix (101414, Quantabio, Beverly, MA, USA). Primers used for qRT-PCR were listed in Table S1. Briefly, qPCR was performed with QuantiStudio3 (Thermo-Fisher) using SYBR Green Real-Time PCR Master Mix (4309155, Thermo-Fisher) in a 10 μL reaction. The standard amplification protocol was 95 °C for 3 min followed by 40 cycles of 95 °C 15 s, 60 °C for 60 s. The *CsGAPDH* gene was used as an endogenous control. All reactions were performed in triplicate. Relative gene expression and statistical analysis were calculated using the 2^-ΔΔCT^ method(Livak and Schmittgen, 2001). qRT-PCR was repeated twice with similar results.

### Microscopy assay

The infiltration areas of *Xcc*-infiltrated leaves and non-*Xcc*-infiltrated leaves from both the wild type *C. sinensis* cv. Hamlin and the *cslob1* mutant were carefully excised using sterilized blades. These excised portions were fixed using 4% paraformaldehyde for a minimum of 2 hours. Subsequently, the specimens underwent a dehydration process before embedded into paraffin chips. The paraffin-embedded chips were cut using a Leica 2155 microtome. Each section was crafted to a thickness of 8 μm. These thin ribbons were positioned onto glass slides and incubated at 37□ overnight to ensure proper heat fixation. Following this, a process of dewaxing and rehydration was performed. The slides were subsequently stained using a 0.05% toluidine blue solution for 30 seconds. After staining, they were rinsed in ddH_2_O, subjected to dehydration, and then a drop of mounting medium was added before covering with a coverslip. Once the mounting medium had solidified, photographs of the slides were captured using the Leica LasX software (Leica Biosystems Inc., Lincolnshire, IL, USA). These images were taken under a bright-field microscope (Olympus BX61; Olympus Corporation, Shinjuku City, Tokyo, Japan).

### *Xcc* growth assay

For *Xcc* infiltration assay, the pictures were captured at 9 days after infiltration. For *Xcc* spraying assay, *Xcc* strain 306 was spraying onto wild type *C. sinensis* cv. Hamlin and transgene-free *cslob1* mutants at the concentration of 5 × 10^8^ cfu/mL. and the leaves were punctured with syringes, creating 8 wounds/leave before spray. After spraying, the plants were covered with plastic bag to maintain the humidity for 24 hours. Images were taken at 18 days after spray. Leaf discs with a diameter of 0.5 cm were collected from the leaves of the plants and leaf discs were ground in 0.2 mL of sterilized H_2_O. To facilitate the assessment of bacterial concentration, serial dilutions of the grinding suspensions, each amounting to 100 μL, were meticulously spread across NA plates (dilutions ranging from 10^-1^ to 10^-6^). Bacterial colonies were counted after 48 h and the number of CFU (colony-forming units) per cm^2^ of leaf disc was calculated and presented with Prism GraphPad software.

### Quantification of H_2_O_2_ concentration

H_2_O_2_ concentration measurement was conducted (Sels et al., 2008; Liu et al., 2023). Briefly, four 0.5-cm-diameter leaf disks from the same leaf that had been injected with water or *Xcc* (1 × 10^7^ cfu/mL) were pooled and stored in a 1.5 mL tube with 0.5 mL of double-distilled (DD) water. The samples were rotated on a platform at 20 rpm for 30 min, and ddH_2_O was replenished with fresh ddH_2_O. Samples were incubated for an additional 6 h on a rotating platform at 20 rpm. H_2_O_2_ concentration was quantified in the supernatants using the Pierce Quantification Peroxide Assay Kit (23280, Thermo Fisher Scientific, Waltham, MA, USA).

## Data availability

The raw reads of genome resequencing for sweet orange plants generated in this study were deposited in the NCBI Bioproject database under the accession number PRJNA1077621. The reference genome of sweet orange was downloaded from Citrus Pan-genome to Breeding Database [http://citrus.hzau.edu.cn/index.php].

## Results

### Genome editing efficacy via Cas12a/crRNAs RNP using 3 crRNAs

For evaluating the gene-editing efficacy via Cas12a/crRNAs RNP using 3 crRNAs, we targeted the *CsLOB1* gene (canker susceptibility gene) by designing three crRNAs. The three crRNAs were located at the promoter region (crRNA1), exon1 region (crRNA2), and exon2 region (crRNA3) (Figure 1A). LbCas12aU/3 crRNAs RNP mixture was transfected into *C. sinensis* cv. Hamlin protoplast cells via PEG-mediated protoplast transformation (Omar et al., 2016b; Su et al., 2023). The DNA samples of transfected protoplast cells at 3 days post transformation (DPT) were isolated and PCR-amplified. The corresponding PCR products were shorter than wild type Hamlin (Figure 1B), which indicated long deletion in the *CsLOB1* gene caused by the Cas12a/3 crRNAs RNP transformation. The short DNA band (Figure 1B) was cut and subjected to colony PCR and Sanger sequencing. All eight colonies were confirmed to harbor different long deletions (Figure 1C).

**Figure 1.**
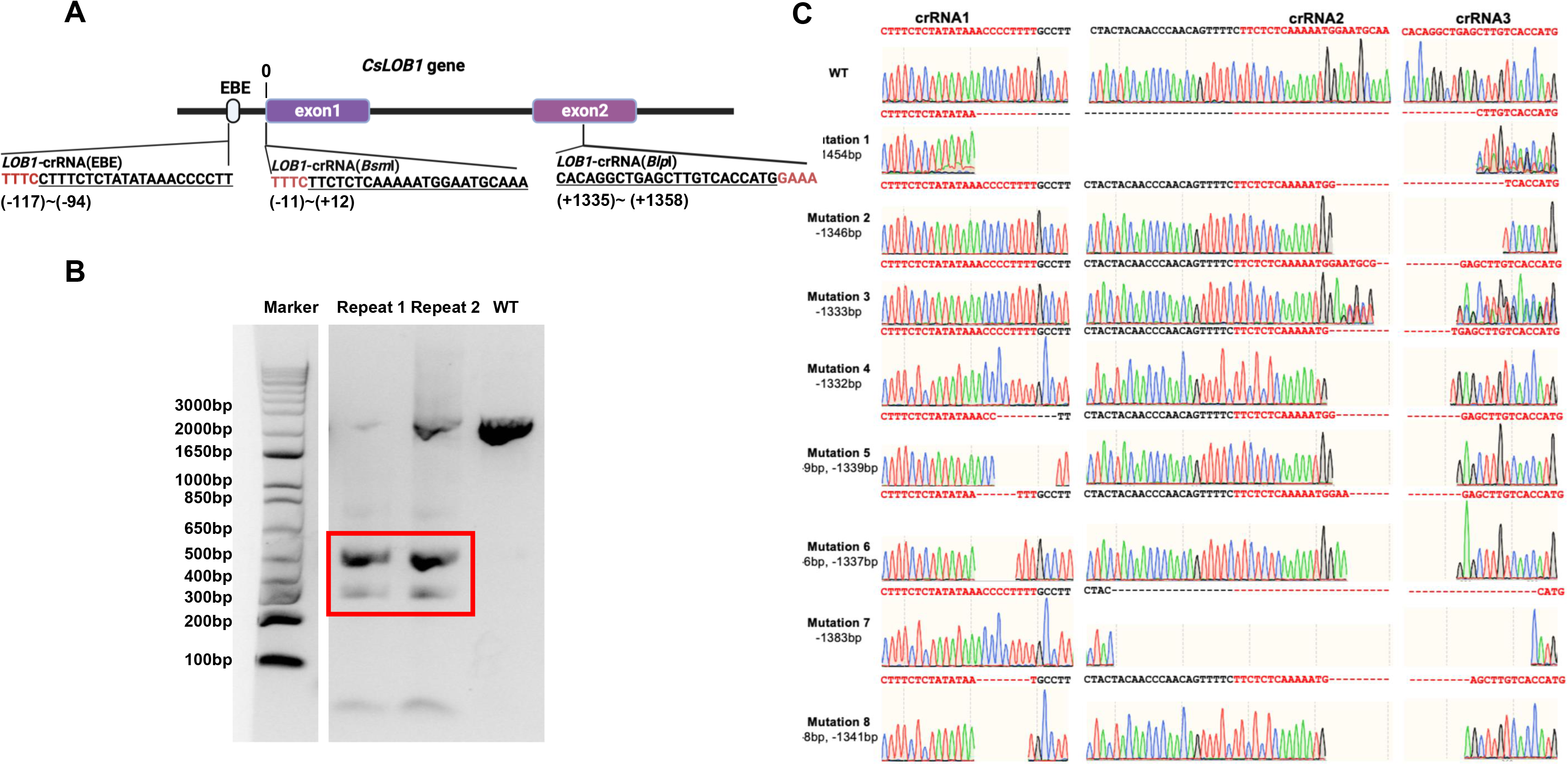
Schematic representation of 3 crRNAs targeting *CsLOB1* gene and evaluation of the crRNA-guided endonuclease activity of Cas12a/3 crRNAs RNP in protoplast at 3 days post transformation (dpt). **A**. The schematic representation of the *CsLOB1* gene and crRNAs. EBE: effector-binding element. Line fragments indicate introns. The PAM (protospacer adjacent motif), either TTTC or GAAA, is highlighted in red. The corresponding crRNAs’ location are labeled below the crRNA sequence. The space between promoter region and the start codon is ‘0’. **B.** Gel picture of *CsLOB1* PCR product amplified from protoplast at 3 days post transformation (dpt). Electrophoresis was performed using a 2% agarose gel. The bands enclosed in the red box correspond to long deletion PCR products, which were subsequently excised, purified, and employed for ligation and colony PCR. **C.** Sequencing results of colonies from PCR products that were excised from the red box region in panel **B**. Among the 8 sequencing reactions, all revealed distinct long deletions associated with crRNAs.

### Transgene-free genome editing of the *CsLOB1* gene in *C. sinensis* cv. Hamlin

Next, we employed Cas12a/crRNAs RNP to generate transgene-free canker-resistant *C. sinensis* cv. Hamlin by editing both promoter region and coding region with 3 crRNAs (Figure 1A). Approximately 9 to 15 months post transformation, we obtained a total of 10 individual regenerated lines from transformed embryogenic citrus protoplasts (Table 1). The 10 regenerated lines exhibited normal growth characteristics akin to the wild type (Figure 2A). Sanger sequencing analysis and whole genome sequencing demonstrated that the 10 lines were mutated in the *CsLOB1* gene (Figure 2B, Figure S1-S20). Among them, 7 lines (#1, 2, 3, 6, 7, 11 and 12) contained biallelic mutations (Table 1). Three lines (#4, 8, and 13) were chimeric (Table 1). Nevertheless, all the edited lines showed 100% mutation rate regardless of being biallelic or chimeric.

**Figure 2.**
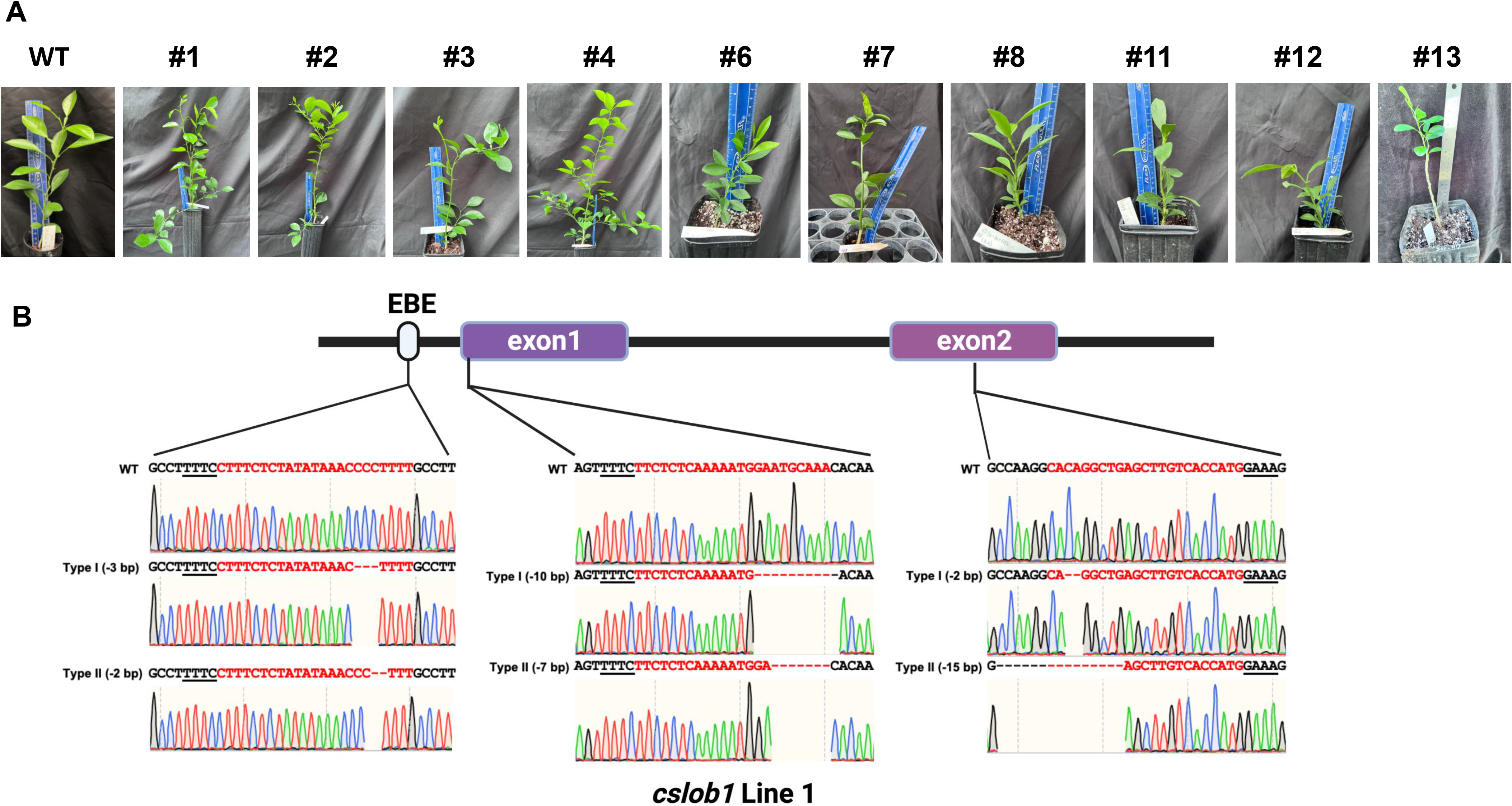
Transgene-free *cslob1* mutants of *C. sinensis* cv. Hamlin and their confirmation via Sanger sequencing. **A**. Transgene-free *cslob1* mutants of *C. sinensis* cv. Hamlin were grafted on Carrizo citrange (*Poncirus trifoliate* × *Citrus sinensis*) rootstock and kept in greenhouse. The genotypes of the mutants were shown. Wild type Hamlin generated from seed was grafted on the same rootstock as a control. The pictures were taken 20 months after potting. **B.** Representative sequencing chromatogram results of transgene-free *cslob1* mutant line 1, which has –3 bp, –10 bp, and –2 bp deletions in type I allele and –2 bp, –7 bp, and –15 bp deletions in type II allele.

**Table 1.** Summary of transgene-free *CsLOB1-*edited *C. sinensis* cv. Hamlin plants generated by LbCas12a-*CsLOB1*(3crRNAs) transformation of embryogenic protoplasts.

| #Line | Mutation type (alleles Type I/II)<br>crRNA1 | Mutation type (alleles Type I/II)<br>crRNA2 | Mutation type (alleles Type I/II)<br>crRNA3 | Genotype |
| --- | --- | --- | --- | --- |
| #1 | -3/-2 | -10/-7 | -2/-15 | Biallelic |
| #2 | -3/-2 | -10/-7 | -2/-15 | Biallelic |
| #3 | -3/-2 | -10/-7 | -2/-15 | Biallelic |
| #4 | -9/(-7,-8) | -1/0 | -7/0 | Chimeric |
| #6 | Inversion/-9 | Inversion/-1344 | Inversion/-1344 | Biallelic |
| #7 | -3/-2 | -10/-7 | -2/-15 | Biallelic |
| #8 | -1460/(-1460,-13) | -1460/(-1460,-9) | -1460/(-1460,-8) | Chimeric |
| #11 | -1454/-1456 | -1454/-1456 | -1454/-1456 | Biallelic |
| #12 | Inversion/-1447 | Inversion/-1447 | Inversion/-1447 | Biallelic |
| #13 | -14/(-1447,inversion) | -1337/(-1447,inversion) | -1337/(-1447,inversion) | Chimeric |

Notably, 5 of the 10 lines contained long deletions, among which 3 lines displayed large inversions (Table 1). Two distinct types of large sequence inversions were identified (Figure 3, Figures S5, S10 and S12). One type (e.g., line 6) involved a large deletion (over 100 bp) between two target sites, followed by inversion (Figure 3A, Figure S5), while the other type (e.g., line 12) displayed a short deletion occurring at three target sites, followed by inversion (Figure 3B, Figure S10).

**Figure 3.**
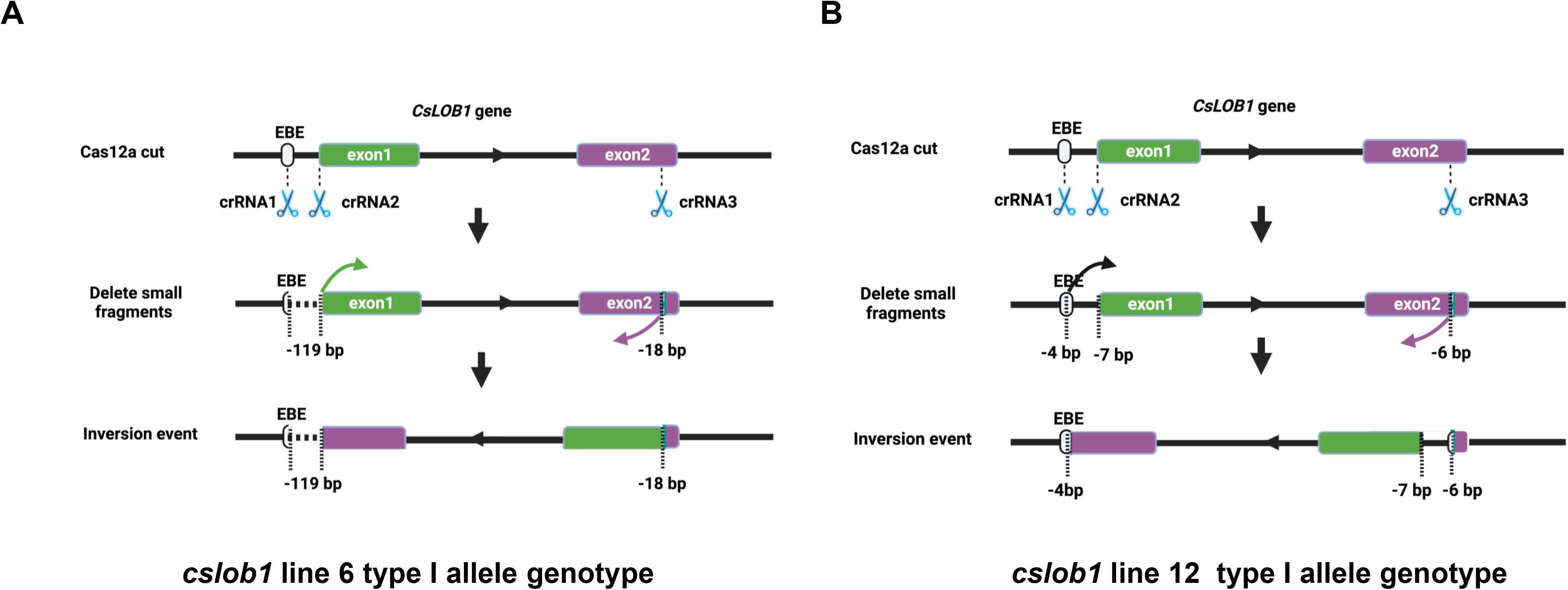
The representative genotypes of inversion events in the *cslob1* mutants via Cas12a/3 crRNAs RNP method. **A**. Type I allele contains inversion in line 6. The three crRNAs resulted in a deletion of 119 bp between the crRNA1 and crRNA2 sites and a deletion of 18 bp at the crRNA3 site. A large fragment as indicated by two arrows was inverted and ligated between the crRNA1 and crRNA3 sites. **B.** Type I allele contains inversion in line 12. The three crRNAs resulted in deletions of 4 bp at the crRNA1 site, 7 bp at the crRNA2 site and 6 bp at the crRNA3 site. A large fragment as indicated by two arrows was inverted and ligated between crRNA1 and crRNA3 sites.

To assess potential off-target mutations, we employed the CRISPR-P v2.0 program (Liu et al., 2017) to identify putative off-target sites with up to 4 nucleotides mismatches with the TTTV PAM site within the citrus genome for the three crRNAs. The analysis revealed zero off-target candidate sites for crRNA1, two candidate sites for crRNA2, and one candidate site for crRNA3 (Table S1). However, off-target mutations were not identified via whole-genome sequencing in all 10 lines.

### Evaluation of *cslob1* mutants

Subsequently, we assessed whether the *cslob1* mutants were resistant to Xcc. Following *Xcc* inoculation for 9 days, the wild type displayed typical canker symptoms, while the *cslob1* mutants exhibited no canker symptoms similar as the non-inoculated leaves (Figures 4A, 4C, & 5). Quantification of *Xcc* titers further corroborated the findings, showing that at 4 dpi and 9 dpi, *Xcc* titers in the wild type were significantly higher than those in the *cslob1* mutants (Figures 4B, & 5). To simulate *Xcc* infection in the natural setting, we conducted *Xcc* inoculation via foliar spray. Typical canker symptoms were observed on wild-type leaves 18 days after spray, whereas no canker symptoms were observed on the *cslob1*mutants (Figure 4D). Moreover, the *Xcc* titer was significantly reduced in the *cslob1* mutants compared to the wild-type at 18 days after spray (Figure 4E).

**Figure 4.**
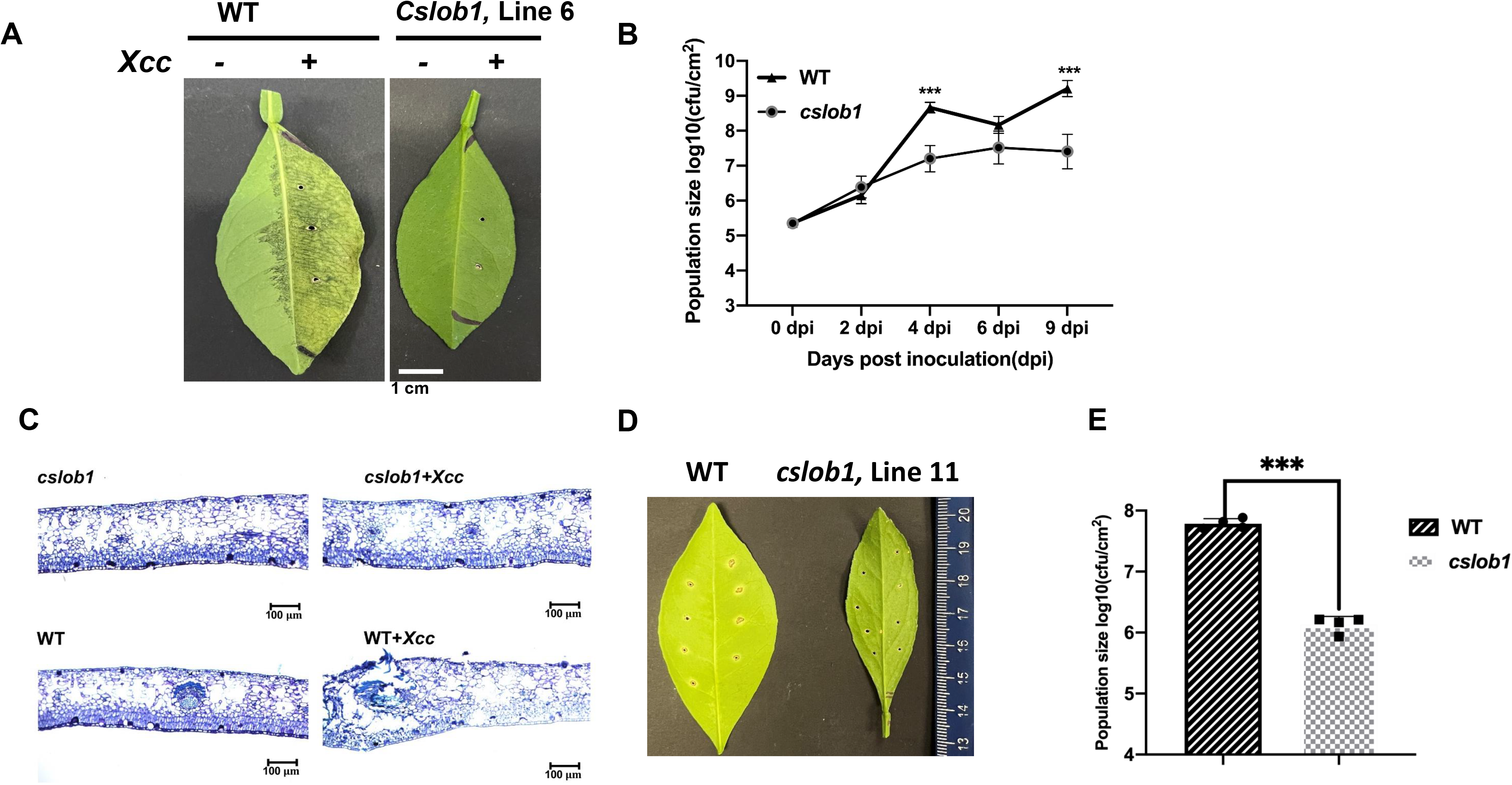
Assessment of disease resistance of the *cslob1* mutant line 6 of *C. sinensis* cv. Hamlin against *Xcc* via injection and foliar spray. **A**. Symptoms of citrus canker on both wild-type and the *cslob1* mutant line 6 of *C. sinensis* cv. Hamlin. The plants were inoculated with *Xcc* through injection using needleless syringes. Inoculation was performed by injecting *Xcc* at a concentration of 1 × 10^7^ CFU/ml into fully expanded young leaves using needleless syringes. The photograph was captured 9 days post inoculation, and a representative image was displayed. The experiment was conducted two times with 3 biological replicates for each experiment with similar results. **B**. *Xcc* growth curve in citrus leaves. Three biological replicates were employed for each experiment and the experiment was repeated two times with similar results. Statistical analysis was conducted using Student’s t-test. Significant differences were indicated by asterisks (P-value<0.05). ‘***’ indicates *P*-value<0.001. (*P*-values at 0 dpi, 2 dpi, 4 dpi, 6 dpi and 9 dpi were 0.8636, 0.2787, 0.0003, 0.0664, and 0.0006, respectively). **C.** Thin cross-section images of **A**. **D** and **E.** Canker symptom and *Xcc* growth of wild-type *C. sinensis* cv. Hamlin and the *cslob1* mutant line 11 after *Xcc* inoculation via spray. Foliar spray was conducted at a concentration of 5×10^8^ cfu/mL and the leaves were punctured with syringes to make 8 wounds/leave before spray. The plants were covered with plastic bag to maintain the humidity for 24 hours after spray. The image was captured at 18 days after spray (**D**) and the *Xcc* titer of **D** was presented €. Student’ s t-test was used for statistical analysis (**E**). ***indicates *P*-value <0.001.

**Figure 5.**
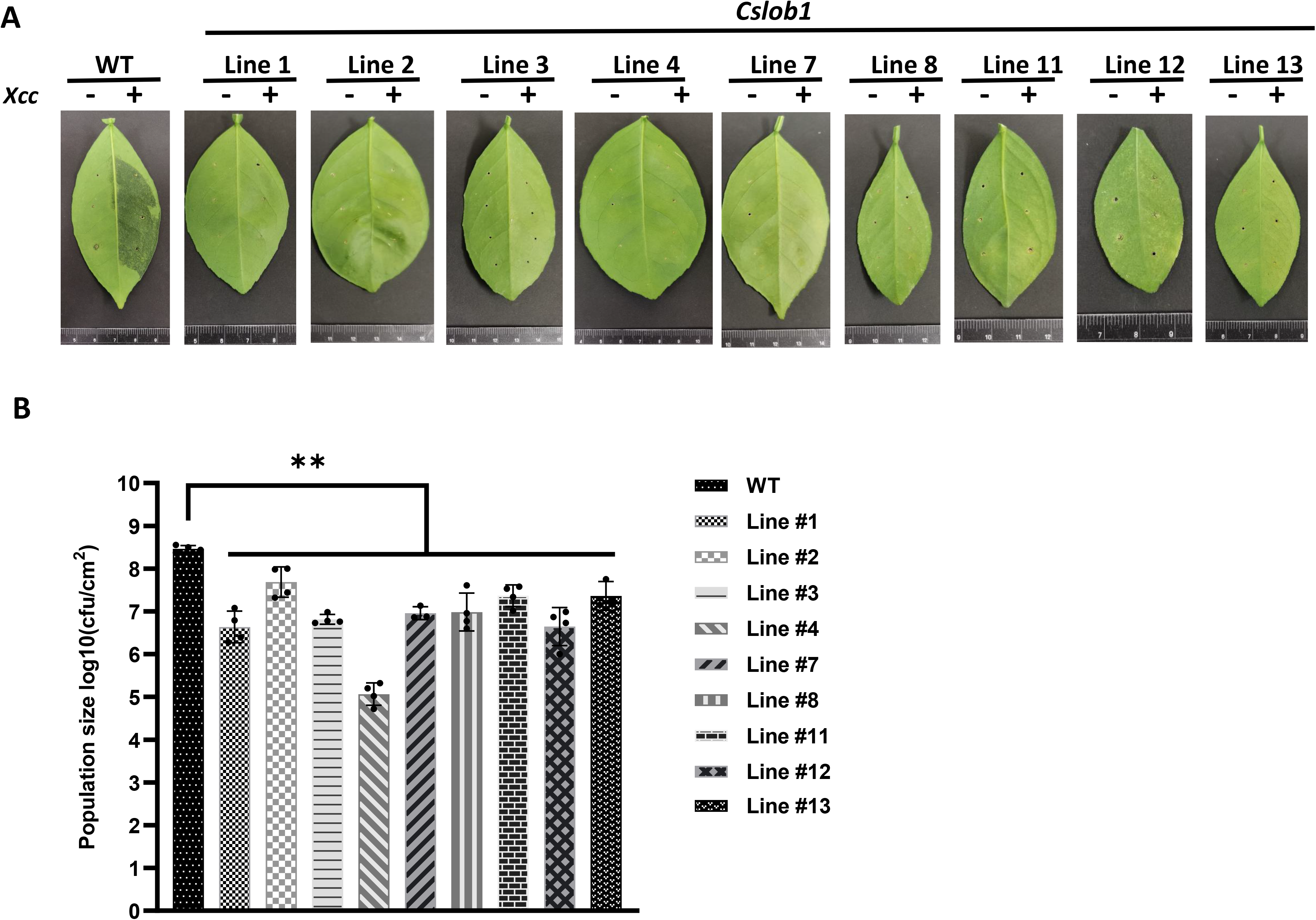
Evaluation of canker resistance of multiple *cslob1* mutants of *C. sinensis* cv. Hamlin via injection. Inoculation was performed by injecting *Xcc* at a concentration of 1 × 10^7^ CFU/ml into fully expanded young leaves using needleless syringes. The photograph was captured 9 days after inoculation (dpi), and a representative image was displayed for each line (**A**). The experiment included three biological replicates, with similar results. **B**. *Xcc* growth in citrus leaves at 9 dpi. Four biological replicates were used, and the mean values ±LJSD (n□=□4) are displayed. The experiments were conducted two times with similar results. Statistical analysis was conducted using Student’s t-test, revealing significant differences indicated by asterisks(P-value<0.05). ‘**’ indicates P-value < 0.01.

Concomitantly, we analyzed the gene expression levels of Cs7g32410 (expansin), Cs6g17190 (RSI-1), and Cs9g17380 (PAR1), known to be up-regulated by *CsLOB1* during *Xcc* infection, 9 days post-*Xcc* inoculation (Zhang et al., 2017c; Duan et al., 2018; Zou et al., 2021). The expression levels of these three genes were significantly reduced in the *cslob1* mutants compared to the wild type plants (Figure 6A). Concurrently, the H_2_O_2_ levels showed no obvious differences in the *cslob1* mutants compared to the wild type (Figure 6B).

**Figure 6.**
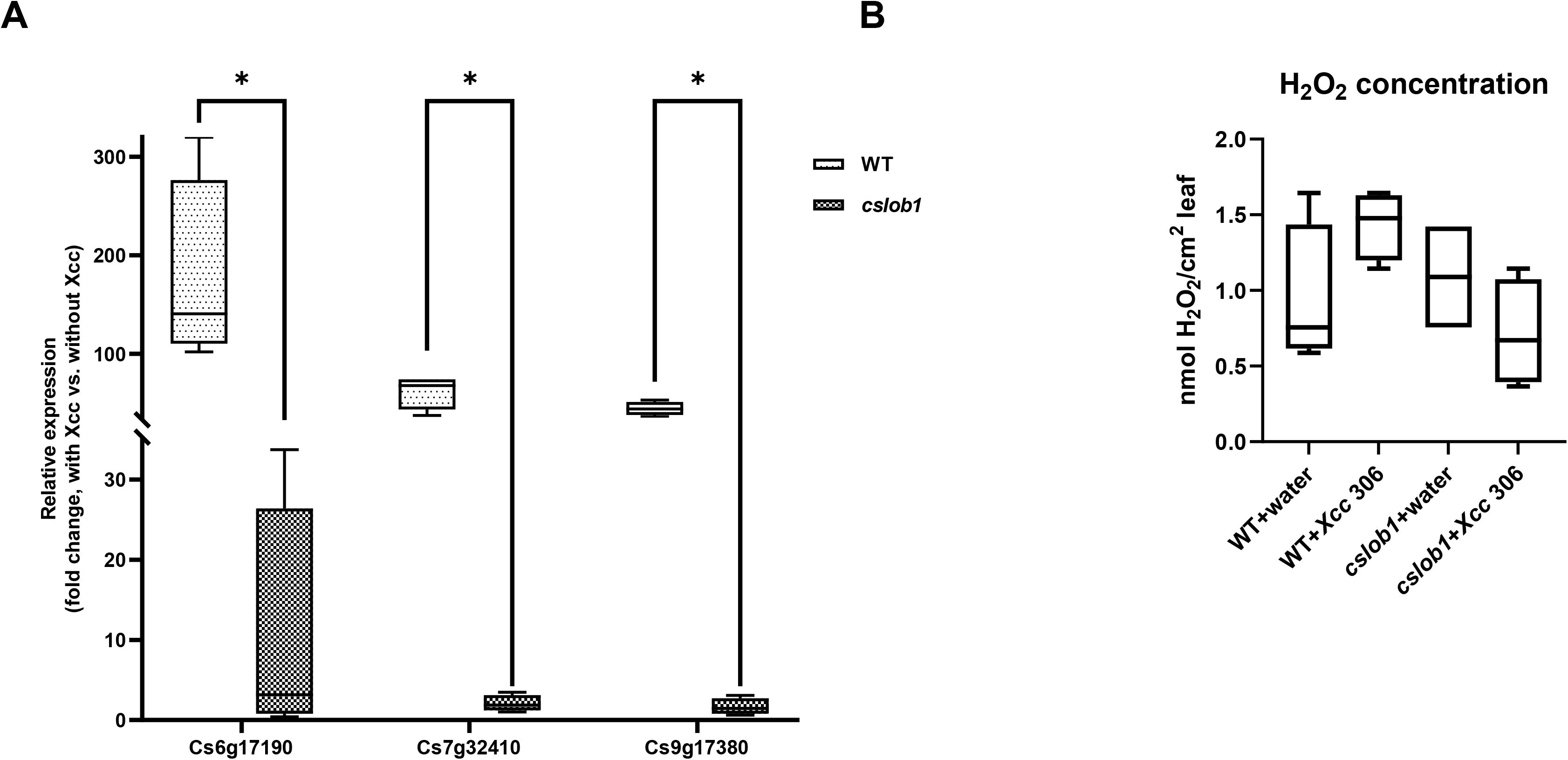
Gene expression and H_2_O_2_ concentration comparison between wild-type and the *cslob1* mutant of *C. sinensis* cv. Hamlin plants in response to *Xcc* inoculation. **A** The expression levels of Cs6g17190, Cs7g32410, and Cs9g17380 genes known to be up-regulated by *CsLOB1* during *Xcc* infection, were examined in both the *cslob1* mutant and wild-type *C. sinensis* cv. Hamlin. This assessment was performed under conditions of *Xcc* inoculation at a concentration of 1 × 10^7^ cfu/mL using syringes. *CsGAPDH*, a housekeeping gene encoding glyceraldehyde-3-phosphate dehydrogenase in citrus, served as the endogenous control. Mean values ± SD (n=4) from four biological replicates were presented. Statistical analysis was conducted using a two-sided Student’s t-test, with single asterisks (*) denoting significant differences (*P* values of Cs6g17190, Cs7g32410, and Cs9g17380 were 0.0412, 0.000447, and 0.0000179, respectively). The experiments were replicated two times with similar results. **B.** Quantification of H_2_O_2_ concentration was performed one day post-inoculation. Each experiment was conducted with four biological replicates. Both the wild type and *cslob1* mutant of *C. sinensis* leaves were subjected to *Xcc* inoculation (1 × 10^7^ cfu/mL) or water, utilizing needleless syringes. The values presented depict means ± SD (n=4). This experiment was repeated twice with similar results. All box plots encompass a median line, where the box represents the interquartile range (IQR), and the whiskers delineate the remainder of the data distribution, excluding outliers. The lower and upper hinges of the boxes correspond to the 25th and 75th percentiles.

## Discussion

In this study, we have demonstrated that transformation of embryogenic protoplasts of citrus with Cas12a/3 crRNAs RNP successfully generates transgene-free biallelic mutants. This is another example of transgene-free genome editing of citrus using this method in addition to a previous report (Su et al. 2023). In the previous report, 38 of 39 of the edited lines were biallelic/homozygous mutants, representing 97.5% mutation rate. In this study, all the edited lines achieved 100% mutation rate with 7 being biallelic mutants whereas 3 being chimeric mutants. It is probable that multiple crRNAs reduce the editing efficacy of individual crRNAs owing to competing for Cas12a but increase the overall editing efficacy. This is consistent with a previous report that higher number of gRNAs leads to reduction of editing efficiency which is likely due to the competition for Cas9 among gRNAs (Xie et al., 2015). Similarly, multiple sgRNAs cause dCas9 bottlenecks in CRISPRi targeted repression by dropping off roughly 1/*N*, where *N* is the number of sgRNAs expressed (Chen et al., 2018a). On the other hand, it was reported that more gRNAs are more efficient than one gRNA in genome editing (Huang et al., 2023). The high genome editing efficacy of transformation of embryogenic protoplasts of citrus with Cas12a/crRNA was also reported by Zhang et al. (Zhang et al., 2022). Despite few reports on generating transgene-free genome-edited citrus through transformation of embryogenic protoplasts with Cas12a/crRNA RNP (Zhang et al., 2022; Su et al., 2023), it has been successfully used in other plants including *Arabidopsis thaliana*, tobacco, lettuce, rice (Woo et al., 2015), grapevine and apple (Malnoy et al., 2016), maize (Svitashev et al., 2016), soybean and tobacco (Kim et al., 2017), wheat (Liang et al., 2017), and cabbage (Murovec et al., 2018), demonstrating enormous potential for its application in genetic improvements of plants.

As expected, transgene-free *LOB1*-edited lines with long deletions were obtained using 3 crRNAs in the RNP complex. Multiple crRNAs induce two or more DSBs in a single chromosome, which is known to mainly lead to deletions (Siebert and Puchta, 2002) and sometimes inversions (Qi et al., 2013; Zhang et al., 2017a). Long deletions were obtained in 5 of the 10 edited lines when 3 crRNAs were used for citrus genome editing, but none of the mutations were long deletions when only one crRNA was used (Su et al., 2023). Additionally, 3 edited lines contained inversions. Importantly, these 3 lines all contained long deletions whereas only two edited lines with long deletions did not contain inversions. This suggests that long deletions are required for occurrence of inversions. The occurrence of inversions suggests that knockin is highly possible when donors are provided in combination with multiple gRNAs, e.g., 3 crRNAs.

Biallelic/homozygous/chimeric mutations were observed for each of the three sites in 4 of the 10 edited lines (#1, #2, #3 and #7) when 3 crRNAs were used. Our data demonstrate that transformation of embryogenic citrus protoplasts with Cas12a/crRNA RNP can be very efficient for multiplex editing. Importantly, this approach avoids the complicated and length process for construction of multiplex vectors or multiple individual vectors (Xie et al., 2015;Čermák et al., 2017; Huang et al., 2020b).

CRISPR genome editing has been widely used for genetic improvements of crops (Zhu et al., 2020). However, most of the genome edited plants have not been commercialized despite improved traits such as yield, quality, and disease resistance. This is because most of the genome-edited plants were transgenic, which require lengthy and expensive deregulation process and are also a concern of consumers. Transgene-free genome editing, on the other hand, can address those issues (Gong et al., 2021; Turnbull et al., 2021; Bhattacharjee et al., 2023). We have previously generated transgene-free canker resistant *C. sinensis* cv. Hamlin plants which were exempted from the regulation of APHIS and is in the process of registration and commercialization (Su et al., 2023). The newly generated transgene-free canker resistant *C. sinensis* cv. Hamlin lines contain longer deletions, indels and inversions and need to go through rigorous field testing for canker resistance and other horticultural traits. As expected, the *CsLOB1* mutations abolish the induction of genes known to be induced by Xcc (Hu et al., 2014; Zhang et al., 2017c; Duan et al., 2018; Zou et al., 2021). The *cslob1* mutants were resistant to citrus canker by reducing growth of Xcc. This reduction probably results from limiting hypertrophy and hyperplasia which are known to provide nutrients to Xcc growth, without affecting ROS production as indicted by H_2_O_2_ (Brunings and Gabriel, 2003).

Overall, Cas12a/3 crRNAs RNP transformation of embryogenic protoplasts of citrus can be used for transgene-free multiplex genome editing with high efficiency. It can be used for deletion of long fragment and has potential for knockin of useful genes.

## Author Contributions

N.W. conceptualized, designed the experiments, and supervised the project. H.S. and Y.W. performed the experiments. A.A.O. and J.W.G. provided citrus suspension culture, J.X. and H.S. performed bioinformatics and statistical analyses. N.W., H.S., J.X., and Y.W. wrote the manuscript with input from all co-authors.

## Acknowledgements

We thank Wang lab members for constructive suggestions and insightful discussions. This project was supported by funding from Florida Citrus Initiative Program, Citrus Research and Development Foundation 18-025, U.S. Department of Agriculture National Institute of Food and Agriculture grants 2023-70029-41280, 2022-70029-38471, 2021-67013-34588, and 2018-70016-27412, FDACS Specialty Crop Block Grant Program AM22SCBPFL1125, and Hatch project [FLA-CRC-005979] to NW.

