## Supplementary Figures and Tables for "Cas12a/3 crRNAs RNP transformation enables transgene-free multiplex genome editing, long deletions, and inversions in citrus chromosome in the T0 generation"

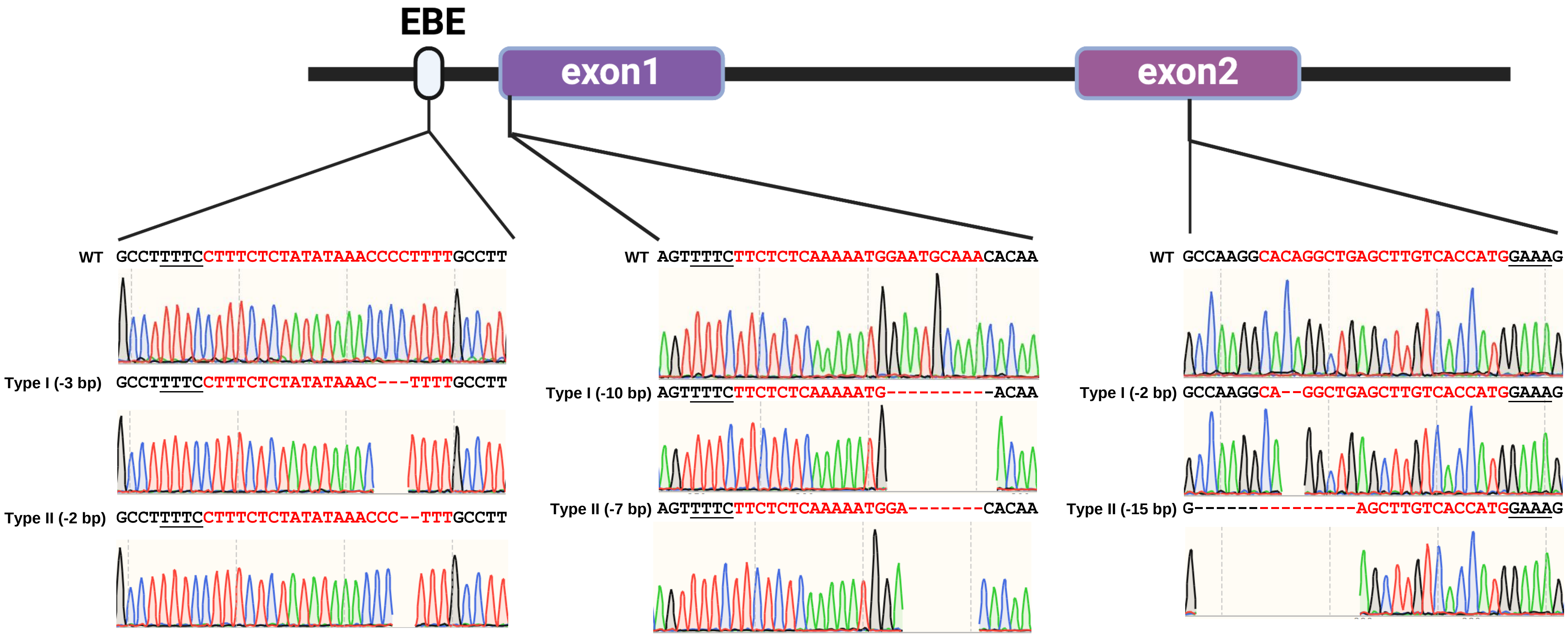


***cslob1* line 2 genotype**

**Figure S1.** Sequencing confirmation of the *CsLOB1* mutation in *C. sinensis* cv. Hamlin line 2 was carried out through PCR amplification and cloning. Representative chromatograms of both alleles of the *CsLOB1* gene were presented for the three crRNAs, revealing short deletions at the targeted crRNA sites. The crRNA sequences were highlighted in red, while the protospacer-adjacent motifs (PAMs, GAAA or TTTC ) were underlined. The symbol "-" was used to denote deletions.

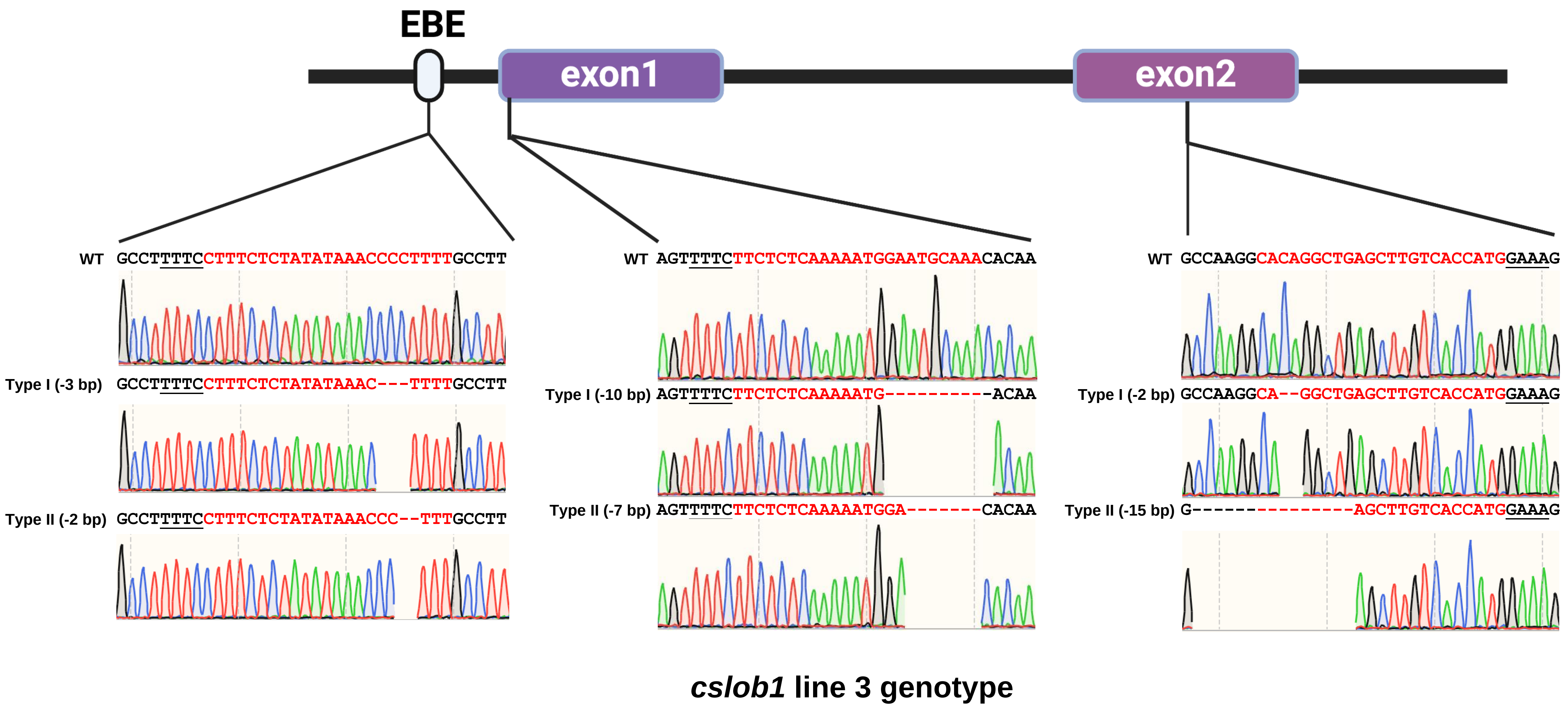


**Figure S2.** Sequencing confirmation of the *CsLOB1* mutation in *C. sinensis* cv. Hamlin line 3 was carried out through PCR amplification and cloning. Representative chromatograms of both alleles of the *CsLOB1* gene were presented for the three crRNAs, revealing short deletions at the targeted crRNA sites. The crRNA sequences were highlighted in red, while the protospacer-adjacent motifs (PAMs, GAAA or TTTC ) were underlined. The symbol "-" was used to denote deletions.

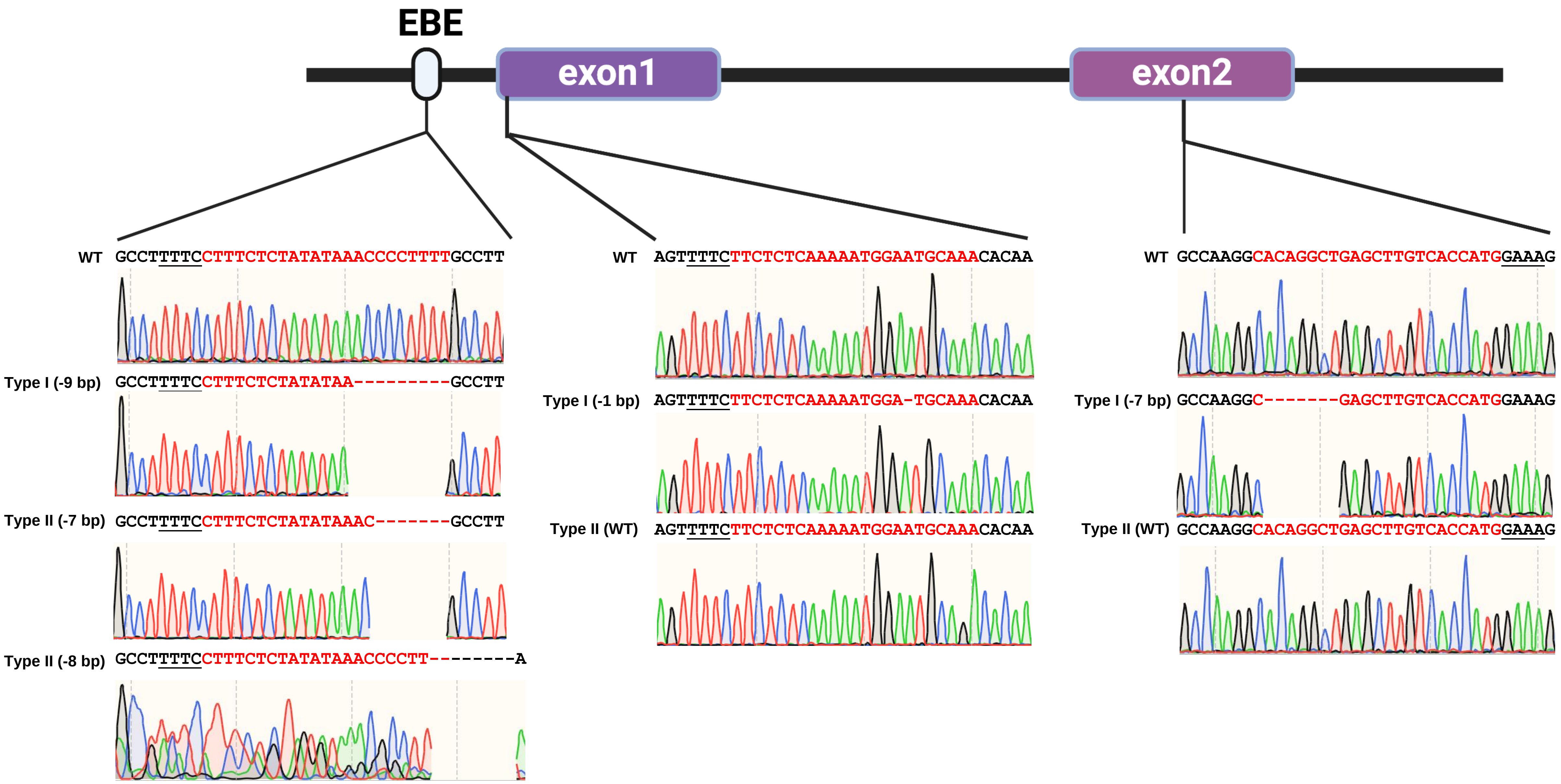


### *cslob1* line 4 genotype

**Figure S3.** Sequencing confirmation of the *CsLOB1* mutation in *C. sinensis* cv. Hamlin line 4 was carried out through PCR amplification and cloning. Representative chromatograms of both alleles of the *CsLOB1* gene were presented for the three crRNAs, revealing small deletions at the targeted crRNA sites. The crRNA sequences were highlighted in red, while the protospacer-adjacent motifs (PAMs, GAAA or TTTC ) were underlined. The symbol "-" was used to denote deletions.

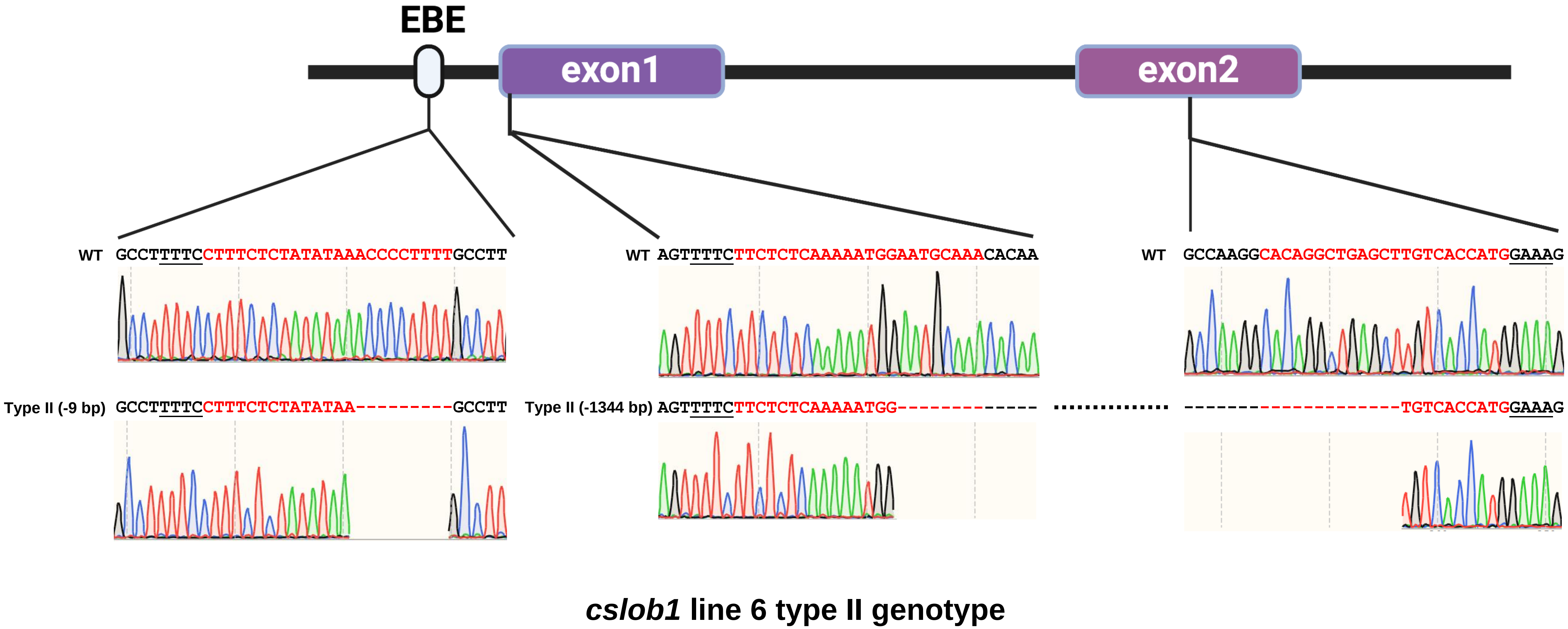


**Figure S4.** Sequencing confirmation of mutation of the type II allele of *CsLOB1* in *C. sinensis* cv. Hamlin line 6 was carried out through PCR amplification and cloning. Representative chromatograms of type II allele of the *CsLOB1* gene were presented for the three crRNAs, revealing deletions at the targeted crRNA sites. The crRNA sequences were highlighted in red, while the protospacer-adjacent motifs (PAMs, GAAA or TTTC ) were underlined. The symbol "-" was used to denote deletions. The dash line indicates long deletion.

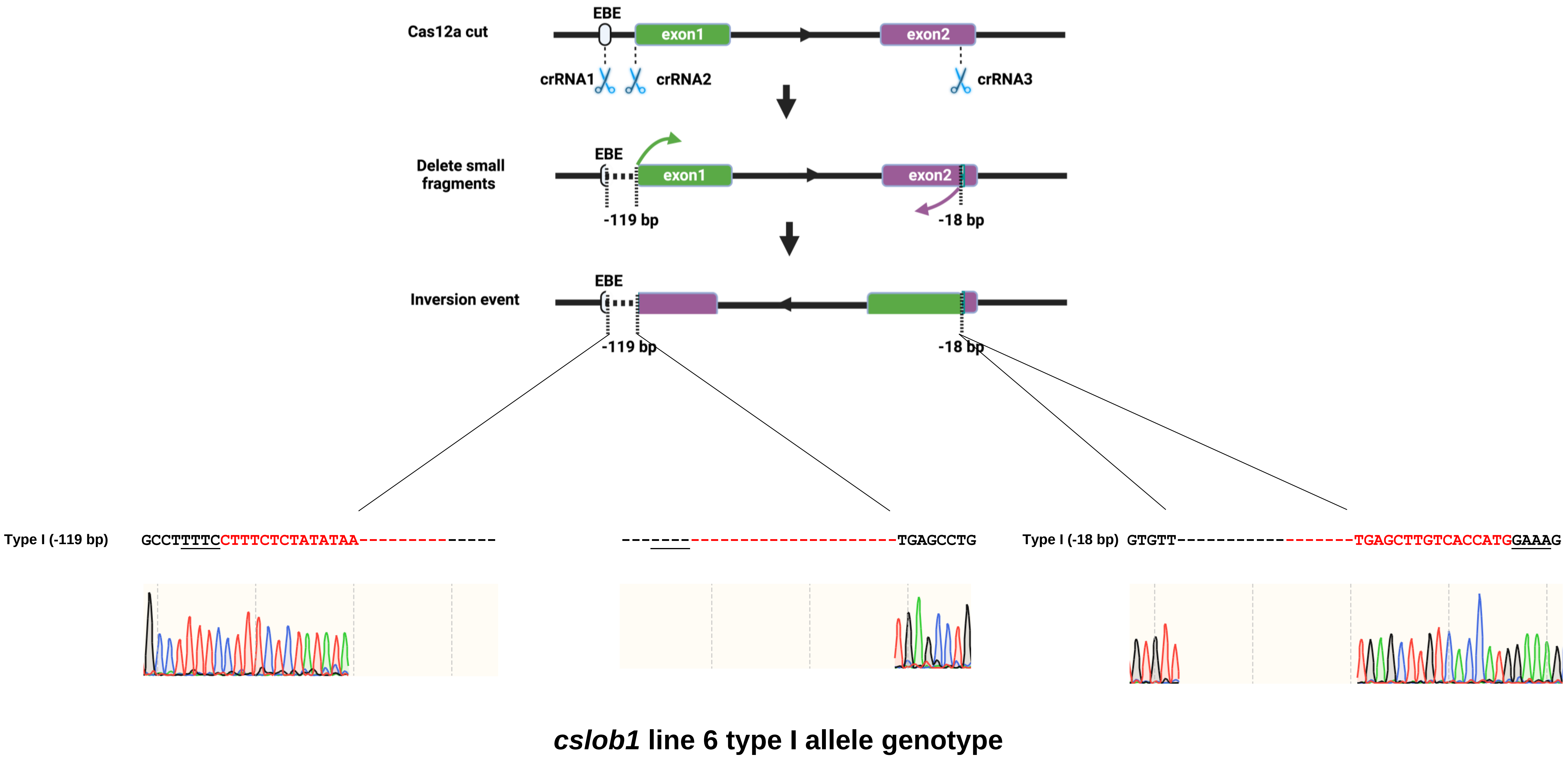


**Figure S5.** Sequencing confirmation of mutation of the type I allele of *CsLOB1* in *C. sinensis* cv. Hamlin line 6 was carried out through PCR amplification and cloning. Representative chromatograms of Type I allele of the *CsLOB1* gene were presented for the three crRNAs, revealing deletions at the targeted crRNA sites and inversion between the two crRNA sites. The crRNA sequences were highlighted in red, while the protospacer-adjacent motifs (PAMs, GAAA or TTTC ) were underlined. The symbol "-" was used to denote deletions.

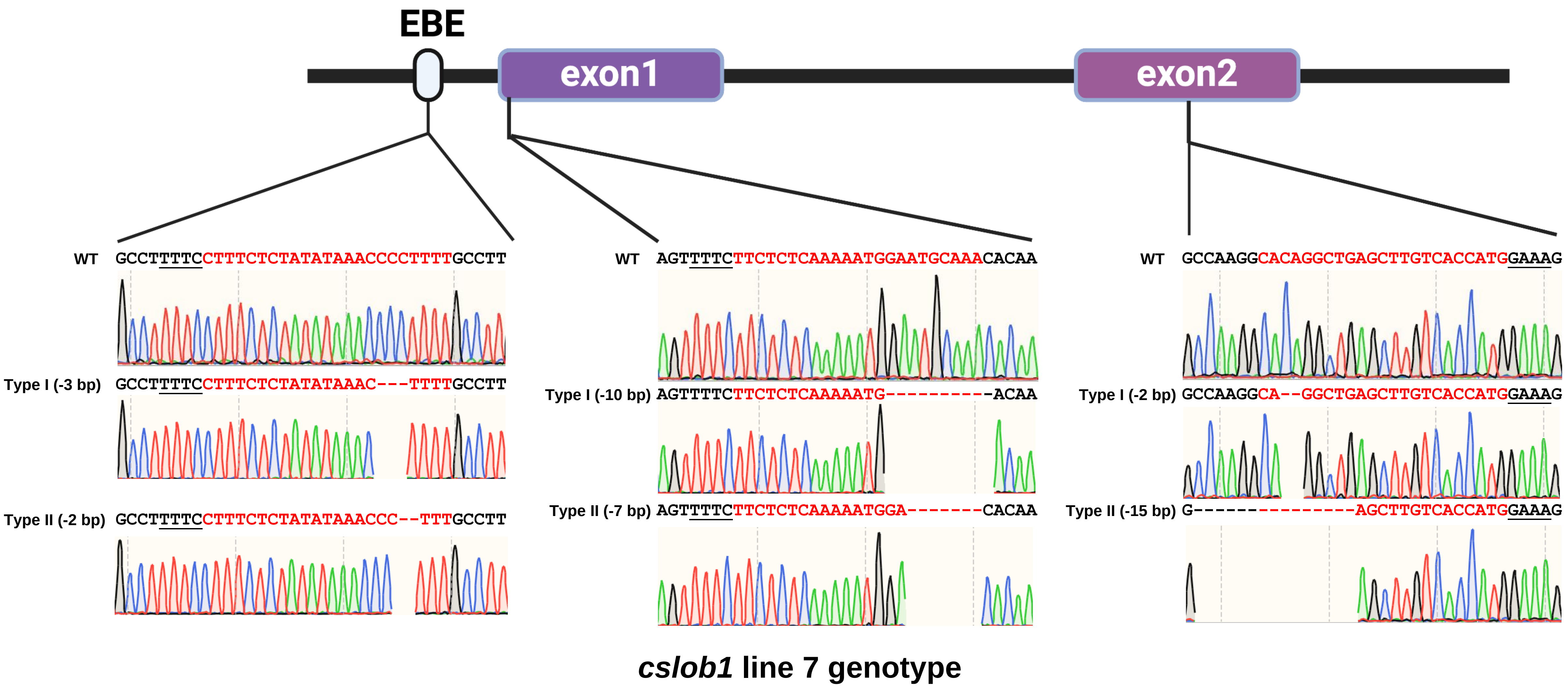


**Figure S6.** Sequencing confirmation of the *CsLOB1* mutation in *C. sinensis* cv. Hamlin line 7 was carried out through PCR amplification and cloning. Representative chromatograms of both alleles of the *CsLOB1* gene were presented for the three crRNAs, revealing small deletions at the targeted crRNA sites. The crRNA sequences were highlighted in red, while the protospacer-adjacent motifs (PAMs, GAAA or TTTC ) were underlined. The symbol "-" was used to denote deletions.

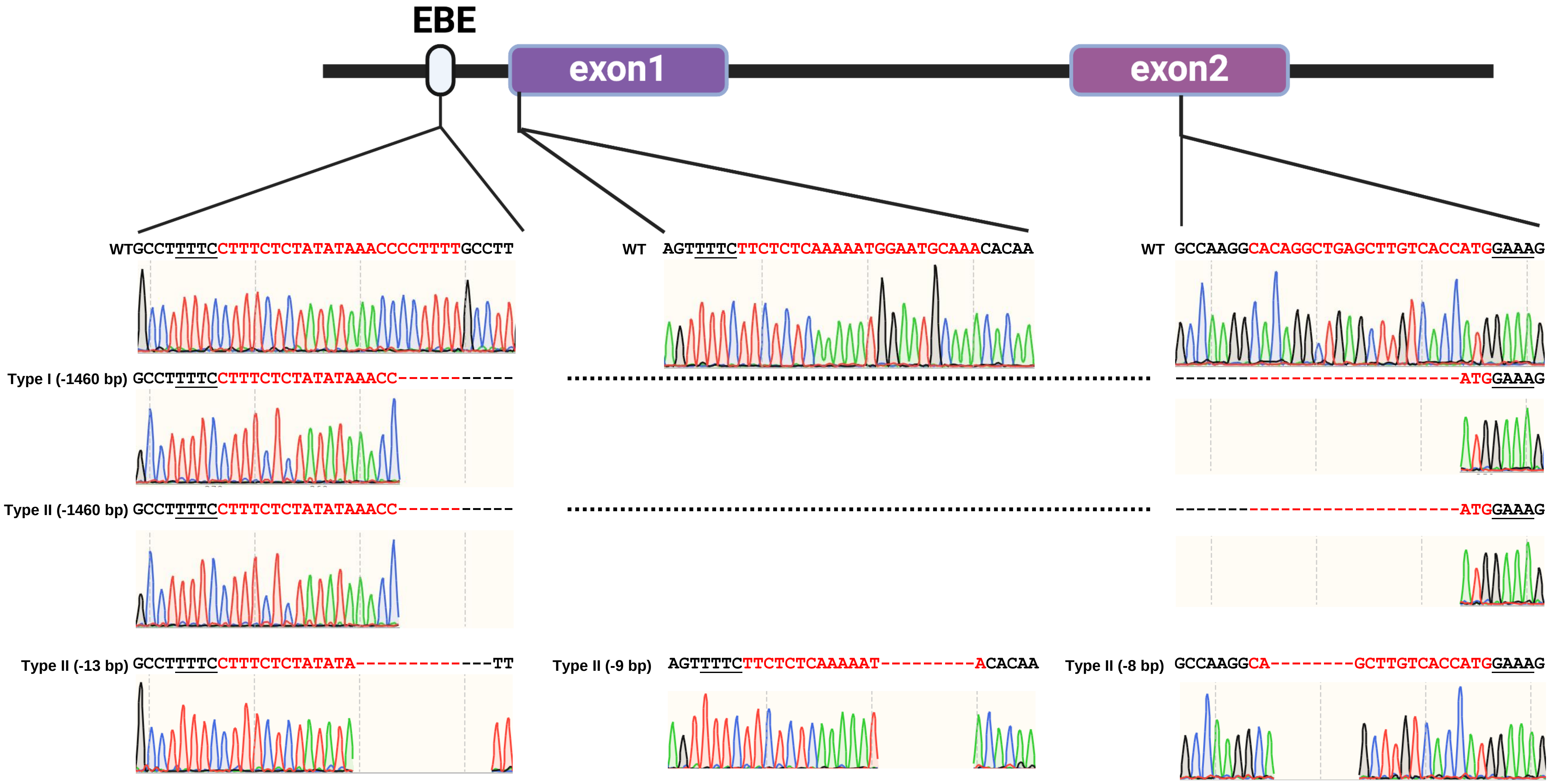


***cslob1* line 8 genotype**

**Figure S7.** Sequencing confirmation of the *CsLOB1* mutation in *C. sinensis* cv. Hamlin line 8 was carried out through PCR amplification and cloning. Representative chromatograms of both alleles of the *CsLOB1* gene were presented for the three crRNAs, revealing long deletions in both alleles. The crRNA sequences were highlighted in red, while the protospacer-adjacent motifs (PAMs, GAAA or TTTC ) were underlined. The symbol "-" was used to denote deletions. The dash line indicates long deletion.

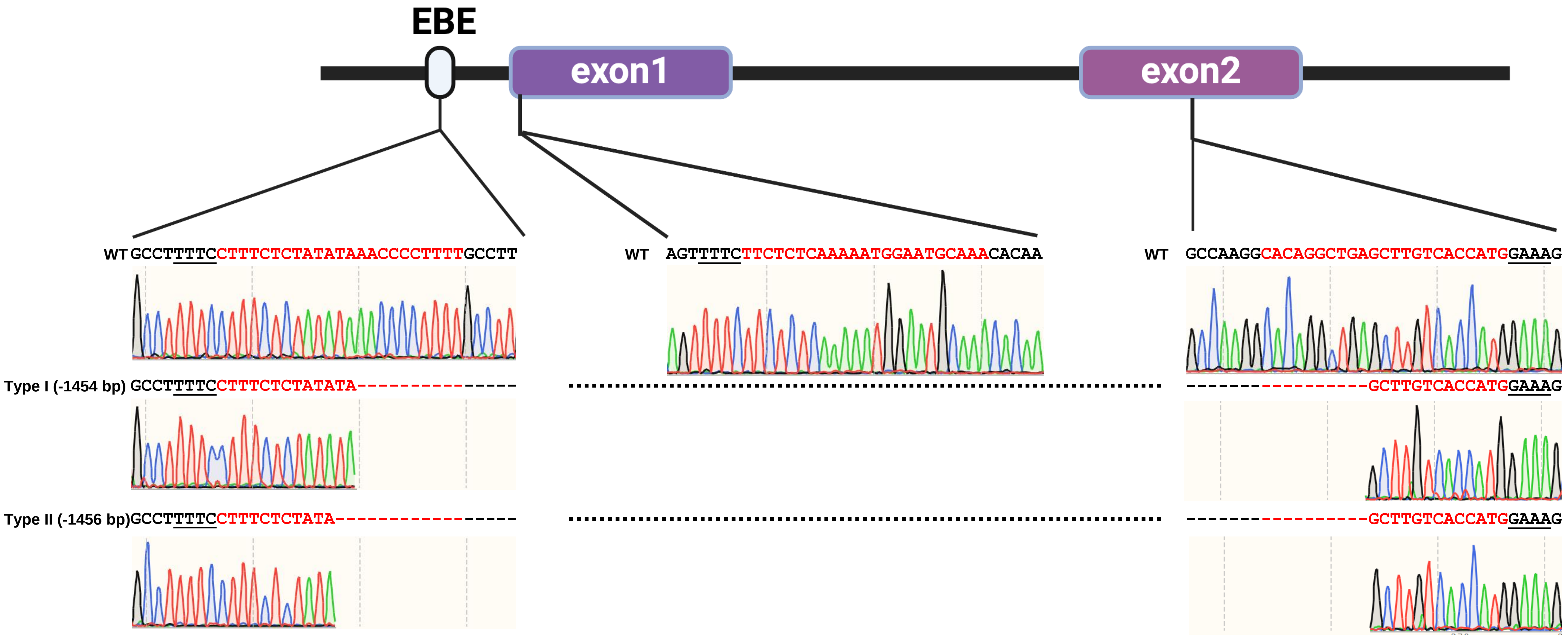


***cslob1* line 11 genotype**

**Figure S8.** Sequencing confirmation of the *CsLOB1* mutation in *C. sinensis* cv. Hamlin line 11 was carried out through PCR amplification and cloning. Representative chromatograms of both alleles of the *CsLOB1* gene were presented for the three crRNAs, revealing small deletions at the targeted crRNA sites. The crRNA sequences were highlighted in red, while the protospacer-adjacent motifs (PAMs, GAAA or TTTC ) were underlined. The symbol "-" was used to denote deletions.

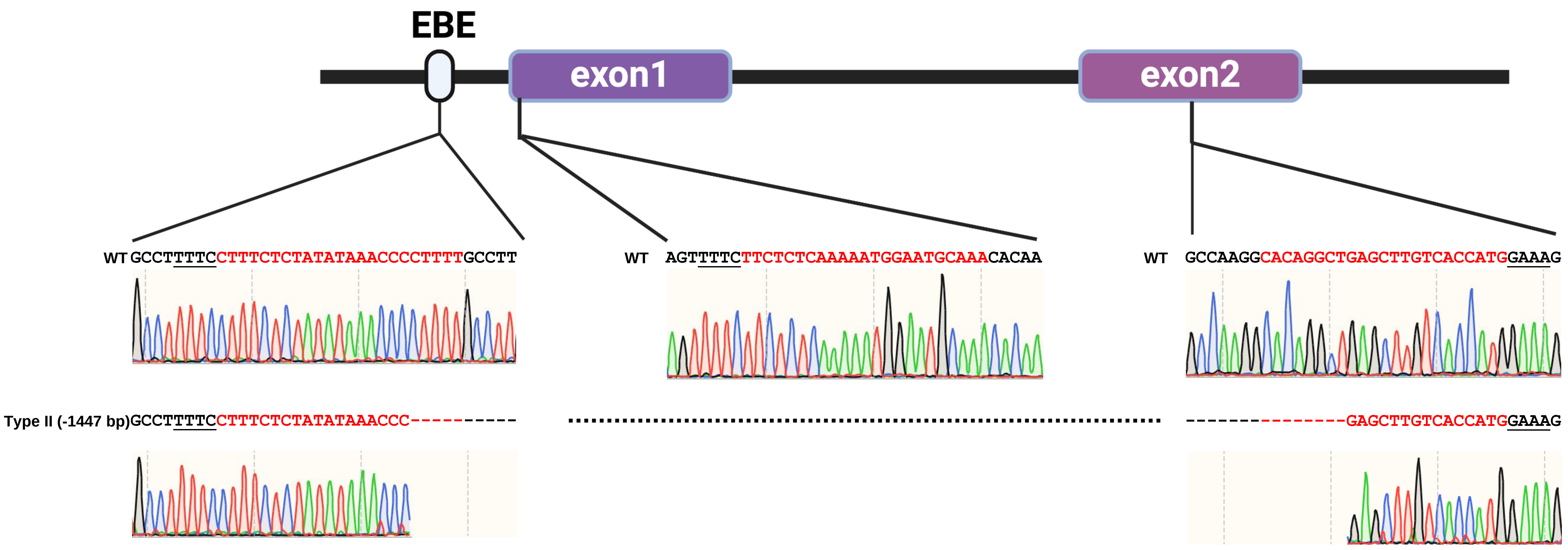


***cslob1* line 12 genotype**

**Figure S9.** Sequencing confirmation of mutation of the type II allele of *CsLOB1* in *C. sinensis* cv. Hamlin line 12 was carried out through PCR amplification and cloning. Representative chromatograms of type II allele of the *CsLOB1* gene were presented for the three crRNAs, revealing deletions at the targeted crRNA sites. The crRNA sequences were highlighted in red, while the protospacer-adjacent motifs (PAMs, GAAA or TTTC ) were underlined. The symbol "-" was used to denote deletions. The dash line indicates long deletion.

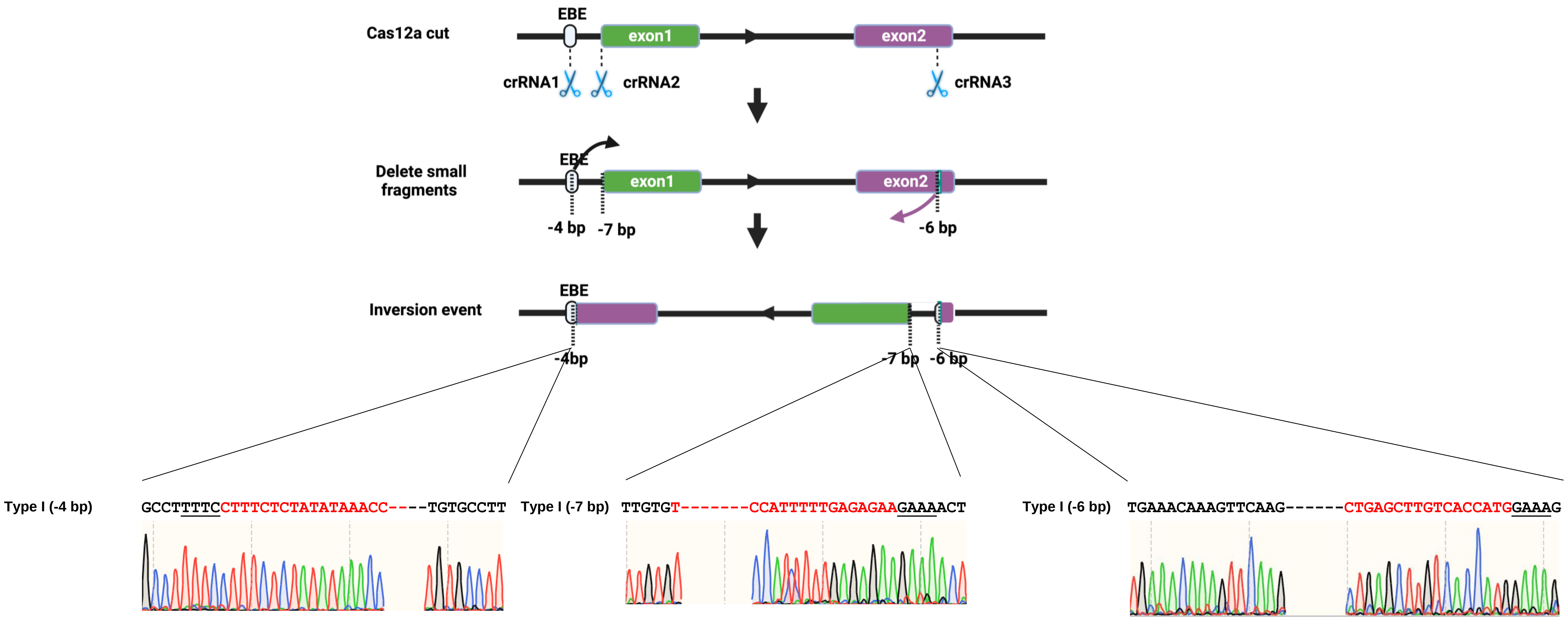


***cslob1* line 12 type I allele genotype**

**Figure S10.** Sequencing confirmation of mutation of the type I allele of *CsLOB1* in *C. sinensis* cv. Hamlin line 12 was carried out through PCR amplification and cloning. Representative chromatograms of Type I allele of the *CsLOB1* gene were presented for the three crRNAs, revealing deletions at the targeted crRNA sites and inversion between the two crRNA sites. The crRNA sequences were highlighted in red, while the protospacer-adjacent motifs (PAMs, GAAA or TTTC ) were underlined. The symbol "-" was used to denote deletions.

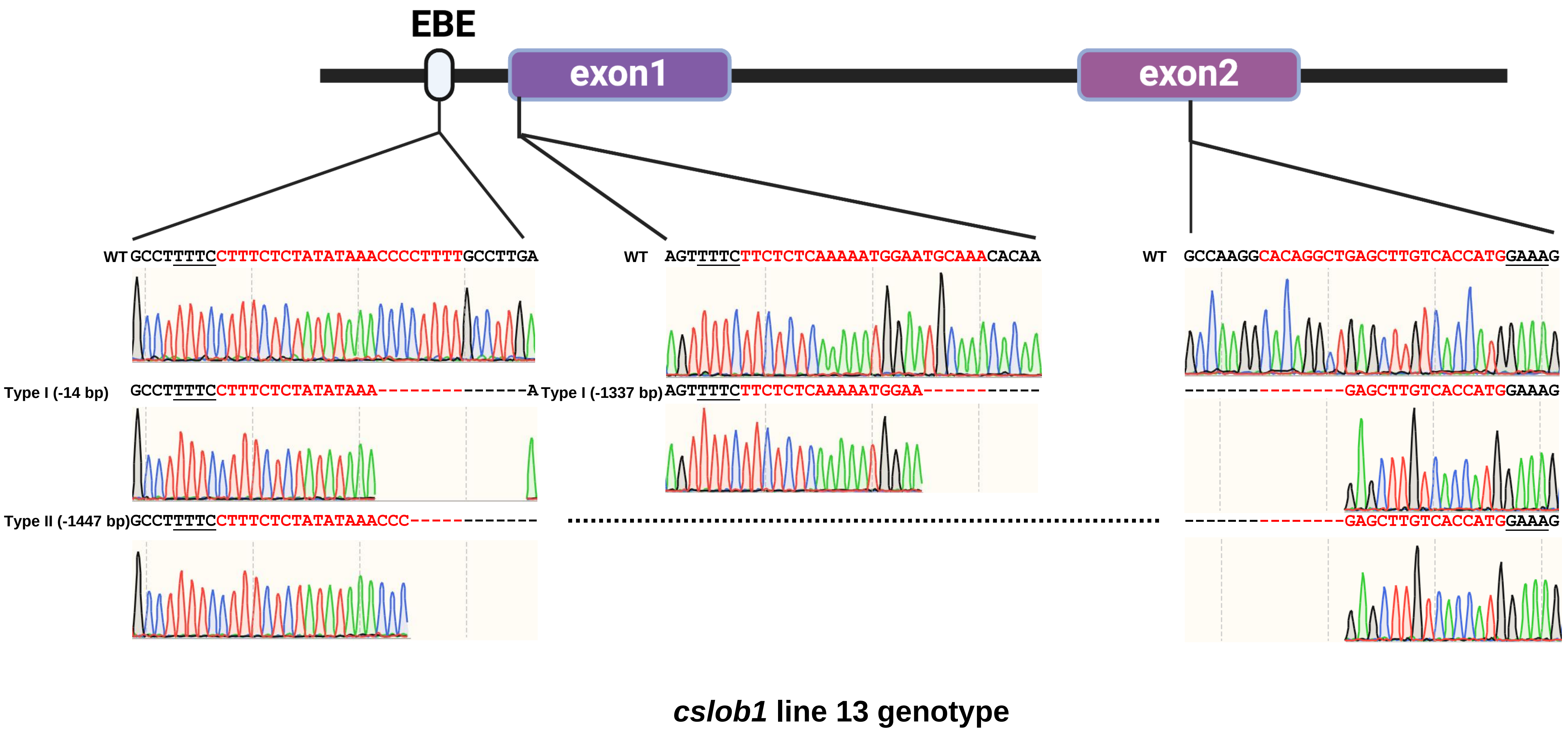


**Figure S11.** Sequencing confirmation of the *CsLOB1* mutation in *C. sinensis* cv. Hamlin line 13 was carried out through PCR amplification and cloning. Representative chromatograms of both alleles of the *CsLOB1* gene were presented for the three crRNAs, revealing small deletions at the targeted crRNA sites. The crRNA sequences were highlighted in red, while the protospacer-adjacent motifs (PAMs, GAAA or TTTC ) were underlined. The symbol "-" was used to denote deletions.

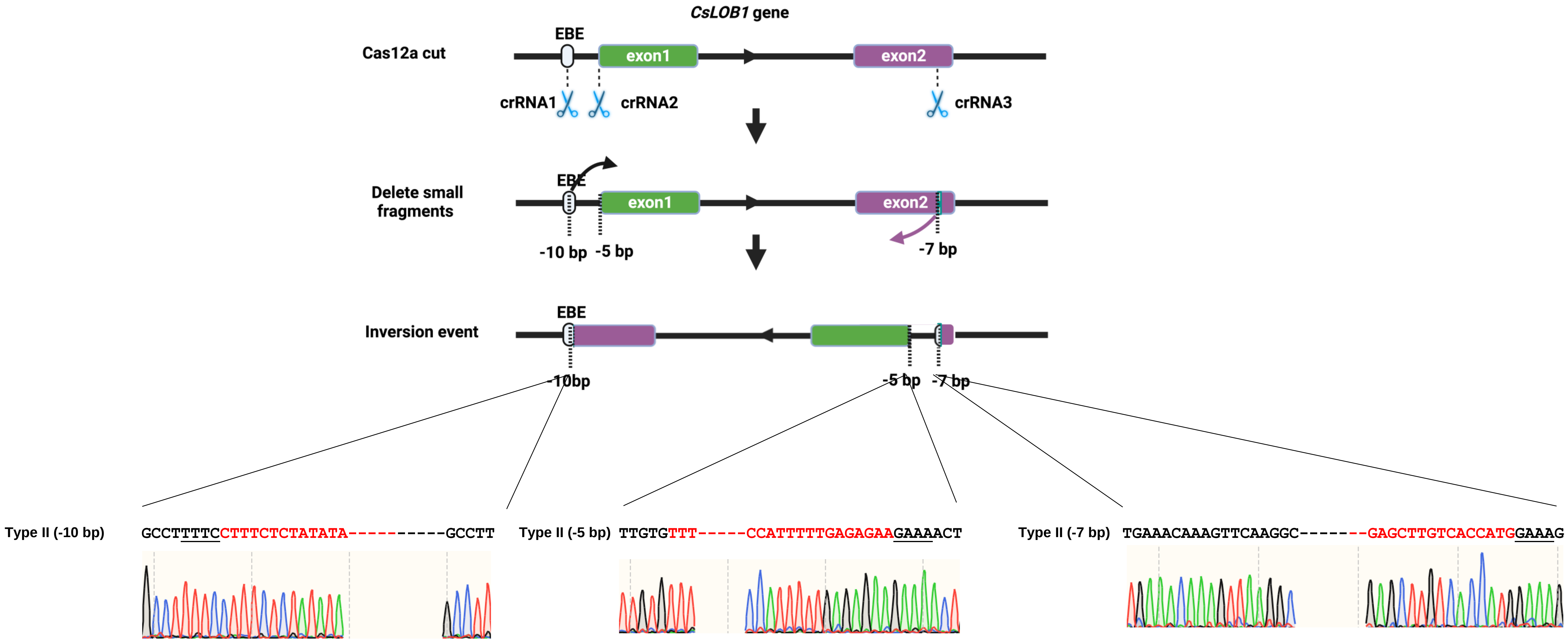


### *cslob1* Line 13 Type II genotype

**Figure S12.** Sequencing confirmation of mutation of the type II allele of *CsLOB1* in *C. sinensis* cv. Hamlin line 13 was carried out through PCR amplification and cloning. Representative chromatograms of Type II allele of the *CsLOB1* gene were presented for the three crRNAs, revealing deletions at the targeted crRNA sites and inversion between the two crRNA sites. The crRNA sequences were highlighted in red, while the protospacer-adjacent motifs (PAMs, GAAA or TTTC ) were underlined. The symbol "-" was used to denote deletions.

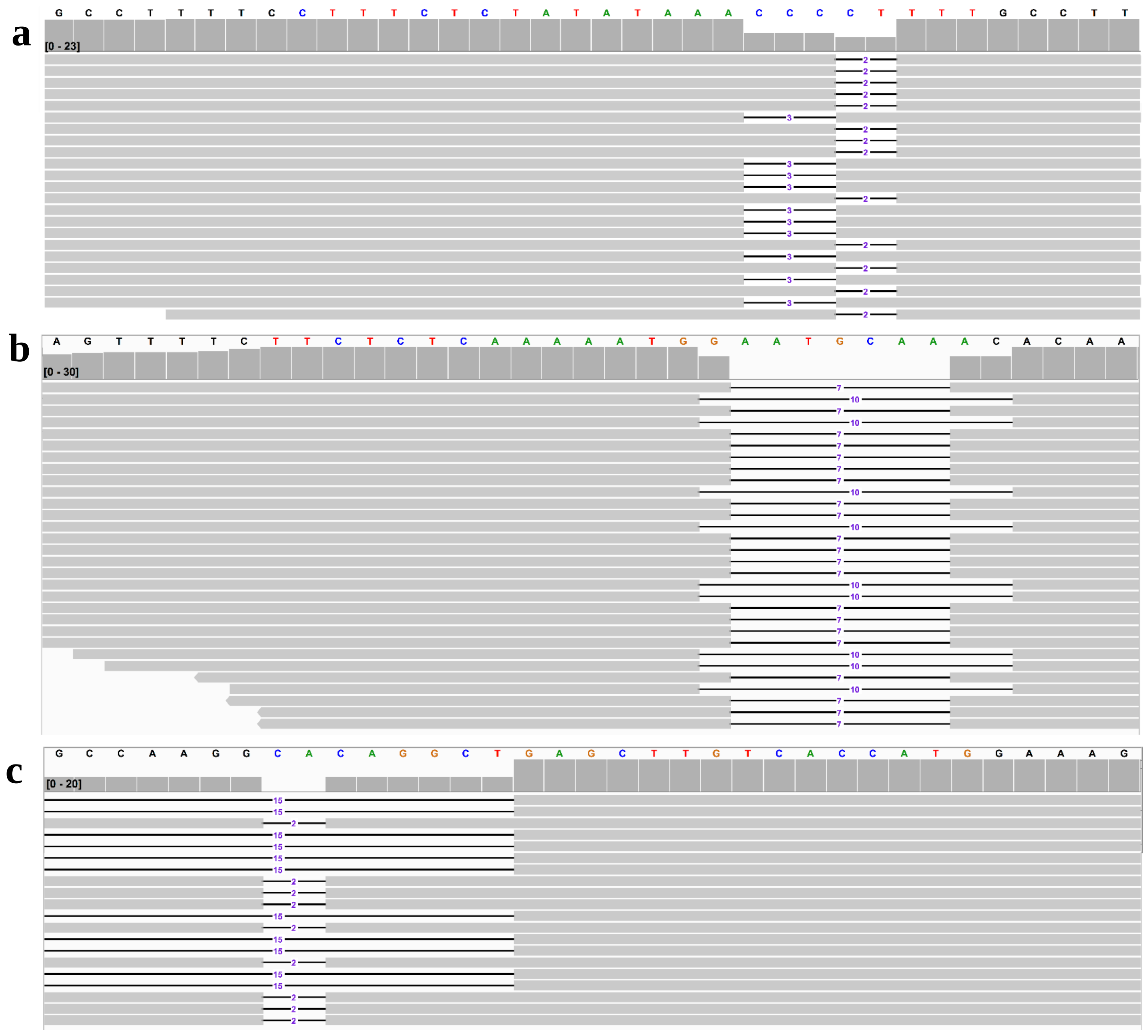


**Figure S13.** Whole genome sequencing identification of the mutations of three target sites of *CsLOB1* gene in line 1. There were two types of deletions of target site of EBE region guided by crRNA1 (a), including type I (-3 bp deletion) and Type II (-2 bp deletion). There were two types of deletions of target site of exon 1 region directed by crRNA2 (b), including type I (-10 bp deletion) and Type II (-7 bp deletion). There were two types of deletions of target site of exon 2 region guided by crRNA3 (c), including type I (-2 bp deletion) and Type II (-15 bp deletion). The bases of target site were highlighted by colors other than black. The mutations were shown by horizontal bar chart. The vertical bar chart showed the sequence depth for each base.

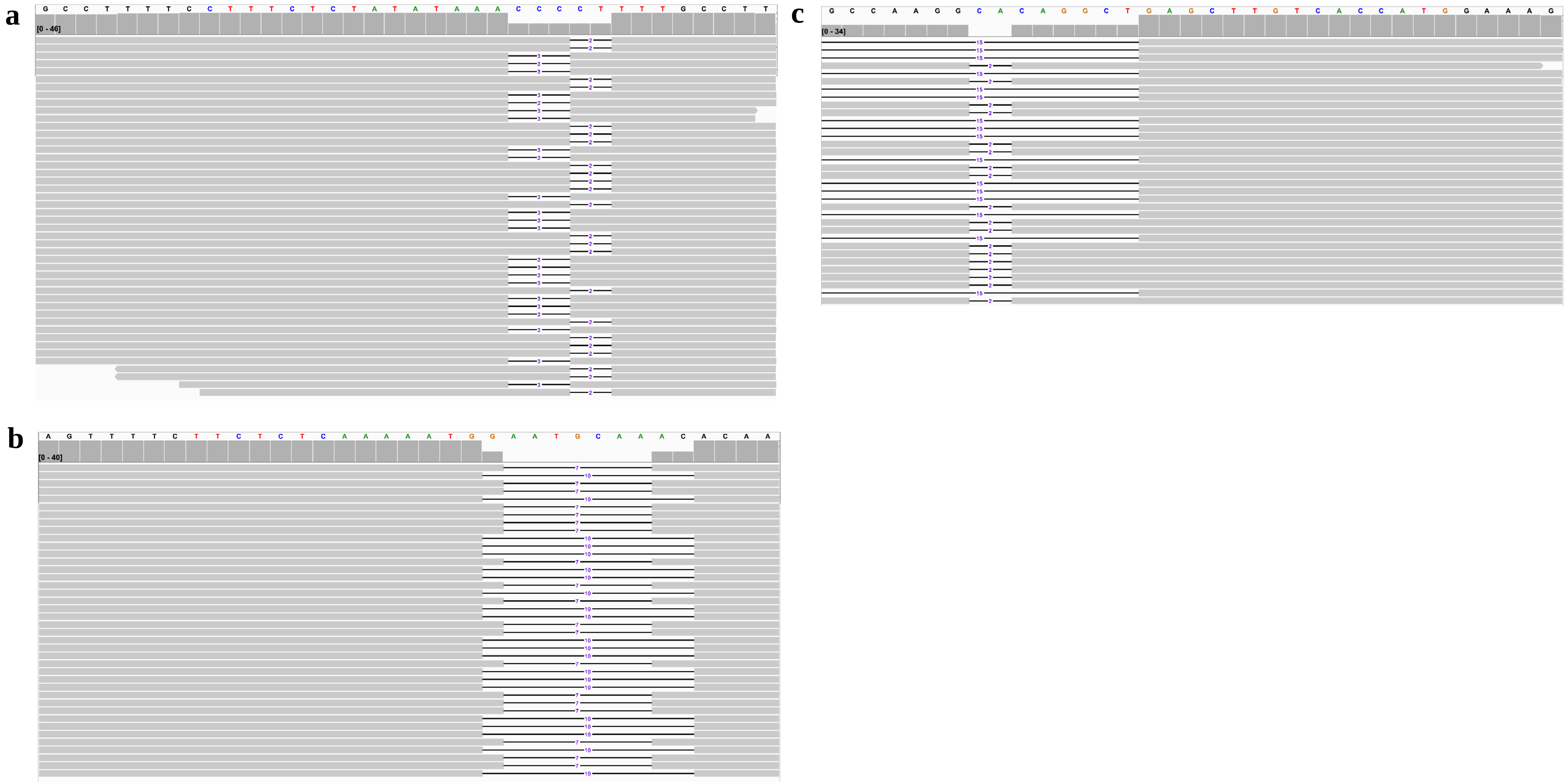


**Figure S14**. Whole genome sequencing identification of the mutations of three target sites of *CsLOB1* gene in line 2. There were two types of deletions of target site of EBE region directed by crRNA1 (a), including type I (-3 bp deletion) and Type II (-2 bp deletion). There were two types of deletions of target site of exon 1 region guided by crRNA2 (b), including type I (-10 bp deletion) and Type II (-7 bp deletion). There were two types of deletions of target site of exon 2 region directed by crRNA3 (c), including type I (-2 bp deletion) and Type II (-15 bp deletion). The bases of target site were highlighted by colors other than black. The mutations were shown by horizontal bar chart. The vertical bar chart showed the sequence depth for each base.

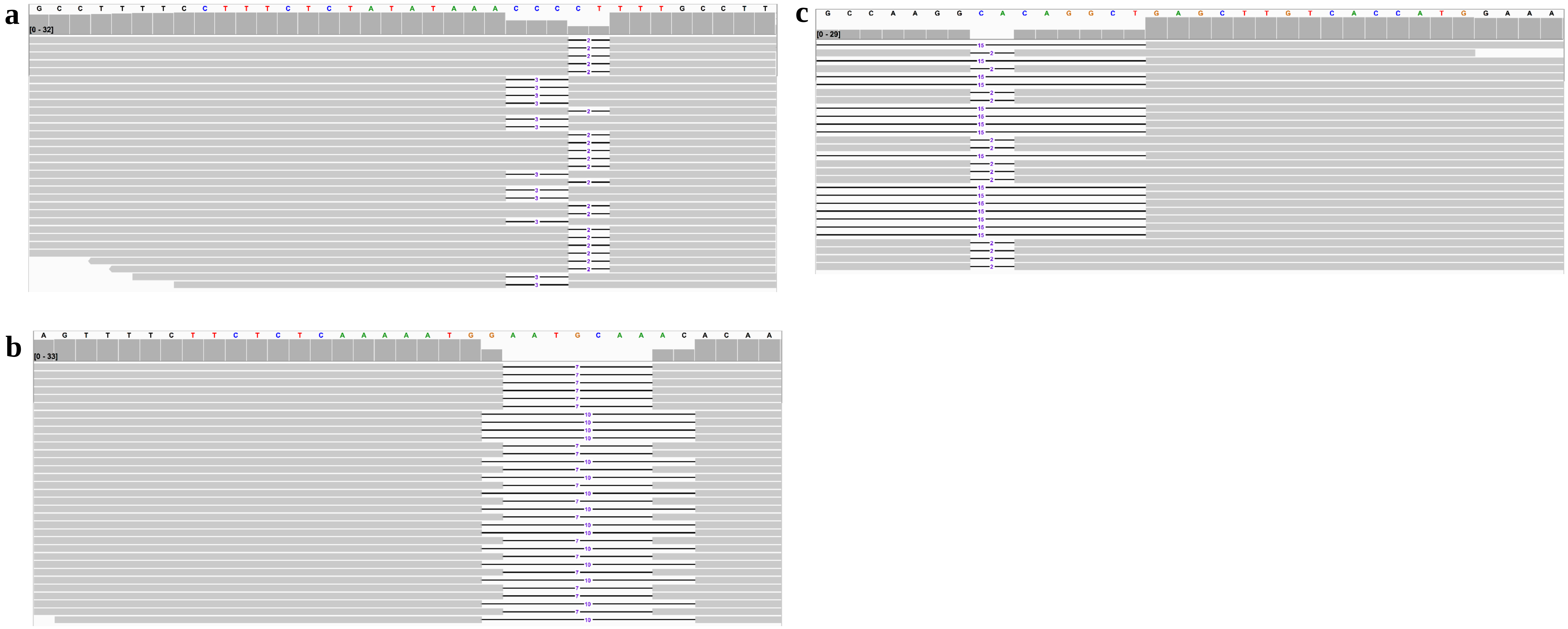


**Figure S15**. Whole genome sequencing identification of the mutations of three target sites of *CsLOB1* gene in line 3. There were two types of deletions of target site of EBE region guided by crRNA1 (a), including type I (-3 bp deletion) and Type II (-2 bp deletion). There were two types of deletions of target site of exon 1 region directed by crRNA2 (b), including type I (-10 bp deletion) and Type II (-7 bp deletion). There were two types of deletions of target site of exon 2 region guided by crRNA3 (c), including type I (-2 bp deletion) and Type II (-15 bp deletion). The bases of target site were highlighted by colors other than black. The mutations were shown by horizontal bar chart. The vertical bar chart showed the sequence depth for each base.

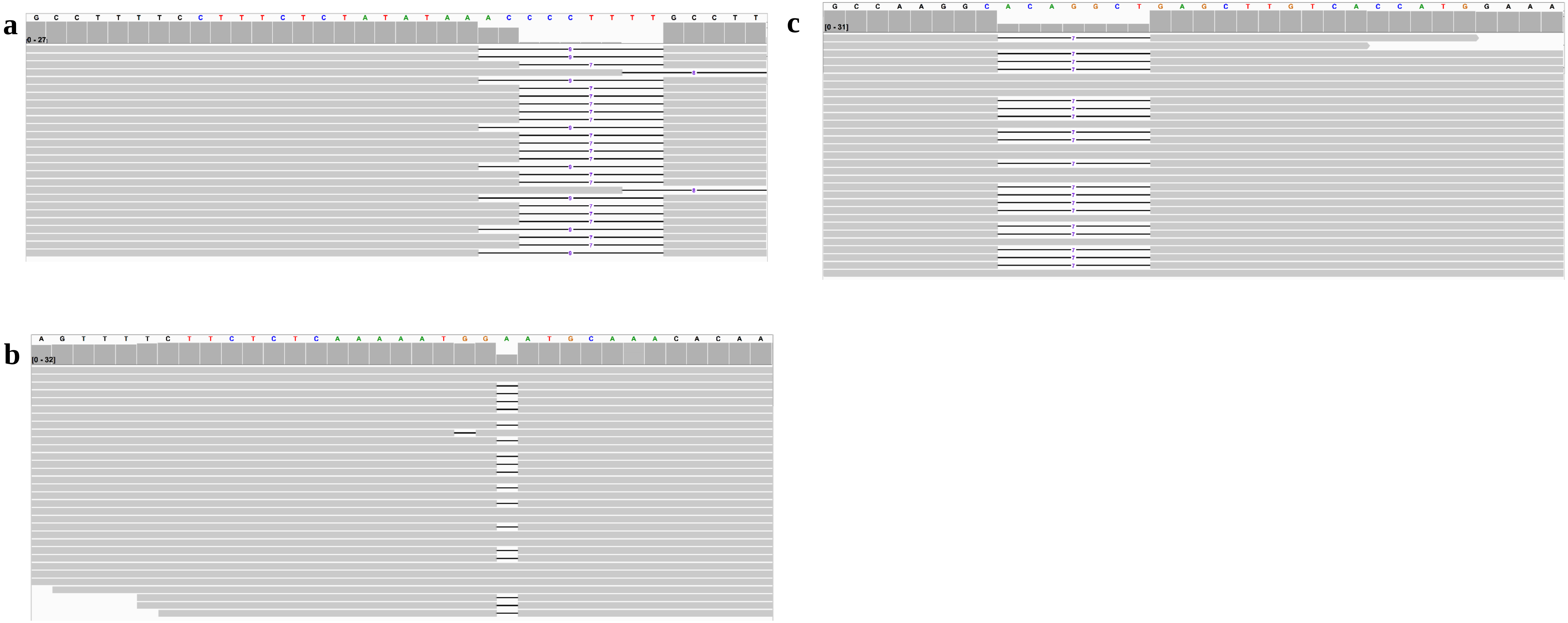


**Figure S16**. Whole genome sequencing identification of the mutations of three target sites of *CsLOB1* gene in line 4. There were three types of deletions of target site of the EBE region directed by crRNA1 (a), including type I (-9 bp deletion) and Type II (-7 bp and -8 bp deletion). There were one type of deletion of target site of exon 1 region (b), including type I (-1 bp deletion), and wild type sequence. There were one type of deletion of target site of exon 2 region (c), including type I (-7 bp deletion), and wild type sequence. The bases of target site were highlighted by colors other than black. The mutations were showed by horizontal bar chart. The vertical bar chart showed the sequence depth for each base.

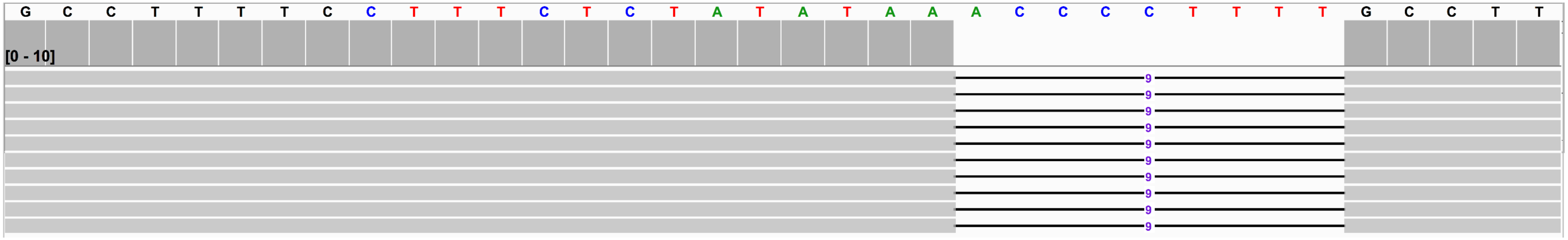


**Figure S17**. Whole genome sequencing identification of the mutations of target site of EBE region of *CsLOB1* gene in line 6. There was one type of deletion of target site of EBE region directed by crRNA1 in type II allele (-9 bp deletion). The bases of target site were highlighted by colors other than black. The mutations were shown by horizontal bar chart. The vertical bar chart showed the sequence depth for each base. Line 6 also contains inversion in allele 1 and 1344 bp deletion in allele 2 which were not shown here.

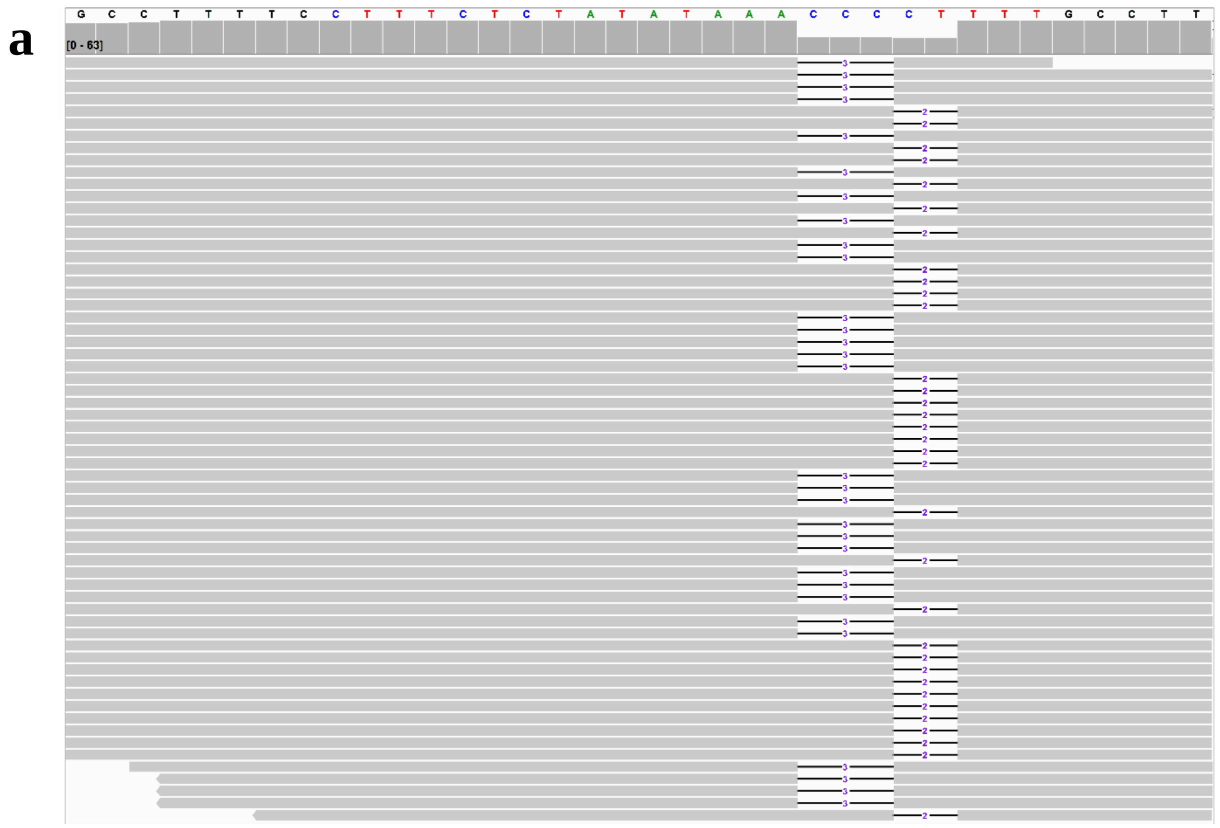

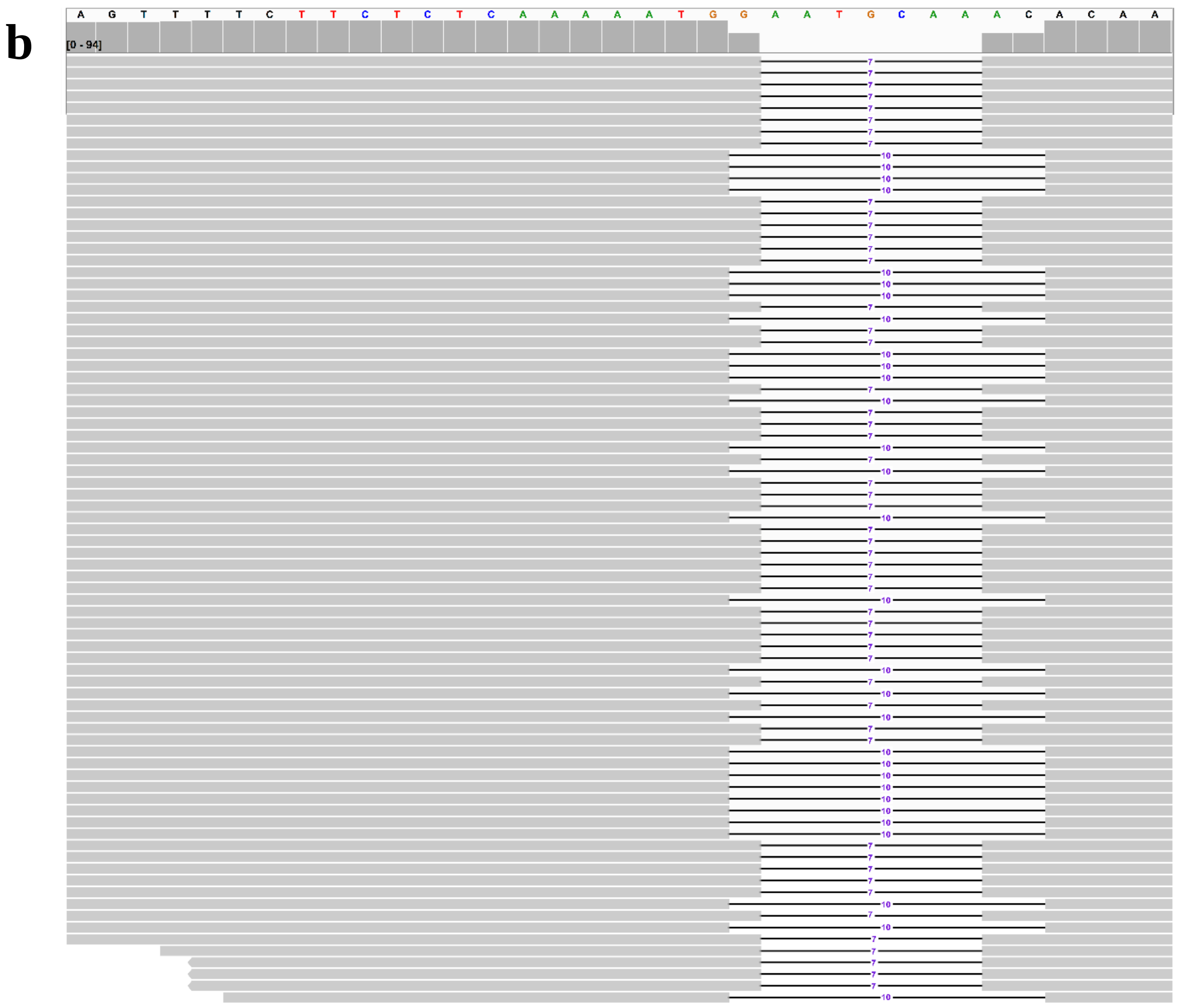

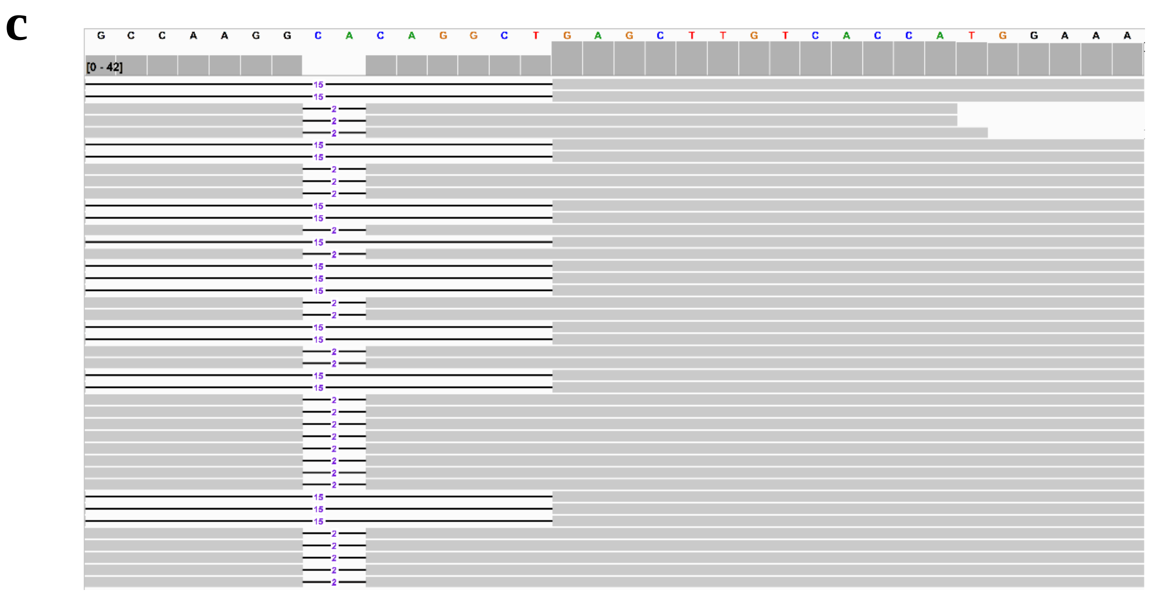

**Figure S18**. Whole genome sequencing identification of the mutations of three target sites of *CsLOB1* gene in line 7. There were two types of deletions of target site of EBE region directed by crRNA 1 (a), including type I (-3 bp deletion) and type II (-2 bp deletion). There were two types of deletions of target site of exon 1 region guided by crRNA2 (b), including type I (-10 bp deletion) and type II (-7 bp deletion). There were two types of deletions of target site of exon 2 region directed by crRNA3 (c), including type I (-2 bp deletion) and type II (-15 bp deletion). The bases of target site were highlighted by colors other than black. The mutations were showed by horizontal bar chart. The vertical bar chart showed the sequence depth for each base.

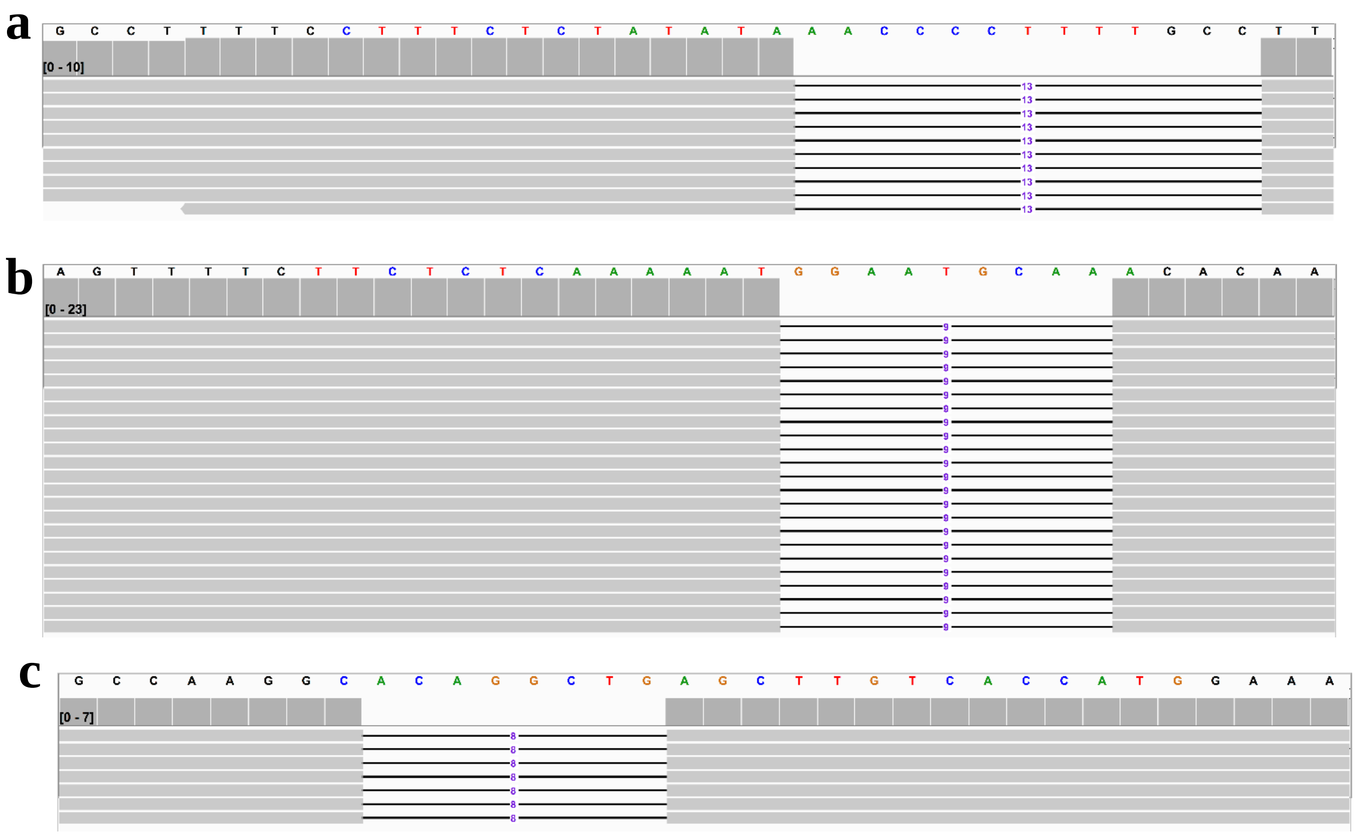


**Figure S19**. Whole genome sequencing identification of the mutations of three target sites of *CsLOB1* gene in line 8. There was one type of deletion of target site of EBE region directed by crRNA1 (a), including type II (-13 bp deletion). There was one type of deletion of target site of exon 1 region guided by crRNA2 (b), including type II (-9 bp deletion). There was one type of deletion of target site of exon 2 region directed by crRNA3 (c), including type II (-8 bp deletion). The bases of target site were highlighted by colors other than black. The mutations were shown by horizontal bar chart. The vertical bar chart showed the sequence depth for each base. Long deletions of 1460 bp were observed in both alleles but not shown here.

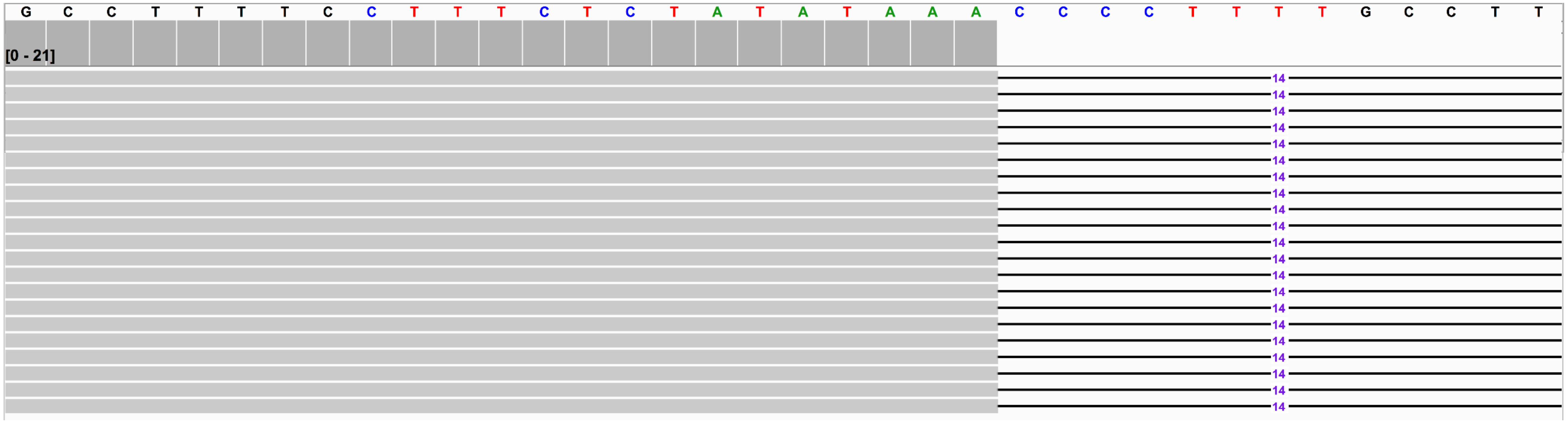


**Figure S20**. Whole genome sequencing identification of the mutations of target site of EBE region of *CsLOB1* gene in line 13. There was one type of deletion of target site of EBE region directed by crRNA1 (a), including type I (-14 bp deletion). The bases of target site were highlighted by colors other than black. The mutations were shown by horizontal bar chart. The vertical bar chart showed the sequence depth for each base. Line 13 also harbors 1337 bp deletion in type I allele and 1447 bp deletion and inversion in type II allele which were not shown here.

Table S1. Primers and crRNAs used in this study

| | |
|---|---|
| Primers for PCR-RE assay | |
| PDS-EXON1-F | ACGATGAGTCCTATTTCCGAGC |
| PDS-EXON1-R | TCCATCCACAATGCCATACACA |
| LOB1-EXON2-F | AGAATCCAGCGCTCAGAGAA |
| LOB1-EXON2-R | TGAATTCCCATCTTGGTTGGGT |
| Primers for mutation detection | |
| F-LOB1-whole | GACATCATCTAGTGGCTCGGTGACAT |
| R-LOB1-whole | TTGATCATGTCCACAGAGGCTCCCAA |
| M13-F | GTAAAACGACGGCCAGTG |
| M13-R | CAGGAAACAGCTATGACC |
| Primers for qRT-PCR | |
| F-Cs7g32410 | GCCTCAGGAACAATGGGAGG |
| R-Cs7g32410 | CCGTGTTAACGCCGTATCCT |
| F-Cs6g17190 | CTCTCGCAGCTCCATTCTGT |
| R-Cs6g17190 | GTCGCCGAACACCGATAAGA |
| F-Cs9g17380 | CTTTGCAGTGGTGGCTCTTG |
| R-Cs9g17380 | TTTGGTCAAGGCTCTCGCAT |
| F-CsGAPDH | GGAAGGTCAAGATCGGAATCAA |
| R-CsGAPDH | CGTCCCTCTGCAAGATGACTCT |
| crRNA sequences | |
| LOB1-crRNA1 | CTTTCTCTATATAAACCCCTTTT |
| LOB1-crRNA2 | TTCTCTCAAAAATGGAATGCAAA |
| LOB1-crRNA3 | CATGGTGACAAGCTCAGCCTGTG |
| Off-target candidates | |
| crRNA2 | TTTCTTCTTCCAAAATTGGAAAGCAAAA |
| | TTTCTTTTTTGAAAAATGGATTGCAAAA |
| crRNA3 | TTTCCAGGTTGCCAACCTCAGCCTGTAC |

Table S2. Summary of genomic sequencing of transgene-free *CsLOB1*-edited *C. sinensis* cv. Hamlin plants

| | Raw data (Gb) | High quality data (Gb) |
|---|---|---|
| #1 | 15.83 | 15.43 |
| #2 | 15.79 | 15.33 |
| #3 | 16.24 | 15.89 |
| #4 | 16.85 | 16.44 |
| #6 | 21.66 | 21.08 |
| #7 | 24.81 | 24.11 |
| #8 | 14.92 | 14.32 |
| #11 | 12.54 | 12.25 |
| #12 | 18.15 | 17.72 |

| | | |
|---|---|---|
| #13 | 20.00 | 19.55 |
